# Cryo-EM structures of PP2A:B55-Eya3 and PP2A:B55-p107 define PP2A:B55 substrate recruitment

**DOI:** 10.1101/2024.09.16.613291

**Authors:** Sathish K.R. Padi, Rachel J. Godek, Wolfgang Peti, Rebecca Page

## Abstract

The phosphoprotein phosphatase (PPP) family of ser/thr phosphatases are responsible for the majority of all ser/thr dephosphorylation in cells. However, unlike their kinase counterpart, they do not achieve specificity via phosphosite recognition sequences, but instead bind substrates and regulators using PPP-specific short linear and/or helical motifs (SLiMs, SHelMs). Protein phosphatase 2A (PP2A) is a highly conserved PPP that regulates cell signaling and is a tumor suppressor. Here, we investigate the mechanisms of substrate and regulator recruitment to the PP2A:B55 holoenzyme to define how substrates and regulators engage B55 and understand, in turn, how these interactions direct phosphosite dephosphorylation. Our cryo-EM structures of PP2A:B55 bound to p107 (substrate) and Eya3 (regulator), coupled with biochemical, biophysical and cell biology assays, show that while B55 associates using a common set of interaction pockets, the mechanisms of substrate and regulator binding can differ substantially. This shows that B55-mediated substrate recruitment is distinct from that observed for PP2A:B56 and other PPPs. It also allowed us to identify the core B55 recruitment motif in Eya3 proteins, a sequence we show is conserved amongst the Eya family. Finally, using NMR-based dephosphorylation assays, we also showed how B55 recruitment directs PP2A:B55 fidelity, via the selective dephosphorylation of specific phosphosites. Because of the key regulatory functions of PP2A:B55 in mitosis and DNA damage repair, these data provide a roadmap for pursuing new avenues to therapeutically target this complex by individually blocking a subset of regulators that use different B55 interaction sites.

## Introduction

Serine/Threonine phosphorylation dependent signaling is essential for the control of most biological processes^1^. Ser/thr kinases, the enzymes that phosphorylate ser/thr residues, typically recognize their substrates using kinase-specific phosphosite recognition sequences, which ensures phosphorylation fidelity and functional signaling events^2^. Ser/thr phosphoprotein phosphatases (PPPs) counteract the action of kinases^1,3,4^. However, a PPP-specific phosphosite recognition sequence has only been identified for PP2B (PP3, Calcineurin [CN])^5^. Indeed, substrate recognition by PPPs, including CN, is predominantly achieved via the recruitment of substrates using PPP-specific short linear motifs (SLiMs; short [4-8 aa] linear stretches of protein sequences that mediate protein-protein interactions) to sites distal from the active site^6–10^. These PPP-specific SLiMs, which are present in either the substrates themselves or scaffolding proteins that recruit substrates, bind directly to preformed SLiM binding pockets on their cognate PPPs. Indeed, PPP-specific SLiMs have been identified for PP1, PP2B/CN, PP2A:B56 and PP4 (PP1, 4-6 SLiMs^8,11–15;^ CN, 2 SLiMs^6,16^; PP2A:B56, 1 SLiM^7,10^; PP4, 1 SLiM^9^). Thus, most PPP-specific inhibitors and substrates use the same interaction pockets for PPP binding, i.e., they compete for the same interaction site(s)^1,17^. Indeed, viruses have exploited this characteristic of PPP-regulator/substrate binding by producing SLiM-containing proteins that inhibit PPP-specific dephosphorylation by binding and blocking PPP regulator/substrate binding sites, rendering them unable to recruit and, in turn, dephosphorylate their specific substrates^6,18,19^. Notably, for PP2B/CN, regulator/substrate binding sites blockage is also used by the immunosuppressant drugs FK-506 and cyclosporin A^6^.

Here we focus on the interaction of substrates and regulatory/scaffolding proteins with PP2A:B55. PP2A is a trimeric holoenzyme comprised of a catalytic subunit, PP2Ac, a scaffolding subunit, PP2Aa, and one of several regulatory subunits, commonly referred to as B subunits each of which adopts a distinct fold (B55, B56, PR72, PR93)^20–22^. The molecular basis of substrate recruitment to B56, which is a heat repeat protein, is well-established. B56 binds substrates and regulators that contain LxxIxE SLiMs, with the binding affinities of distinct SLiMs modulated by phosphorylation and/or adjacent dynamic, charge:charge interactions^7,10,23^. In contrast, much less is known about the mechanism(s) of substrate and regulator recruitment for the most abundant PP2A holoenzyme, PP2A:B55 (B55 is a WD40 domain; **Fig. 1a**)^24,25^. Recently, we determined the structures of two protein inhibitors, phosphorylated-ARPP19 (ARPP19) and FAM122A, bound to PP2A:B55^26^. The inhibitor interactions were extensive, leading to the identification of multiple surfaces on B55 whose shallow pockets collectively mediate tight inhibitor binding, including the B55 platform (pockets P1-P6; ARPP19 and FAM122A), the B55 wall (P7-P9, ARPP19 and FAM122A), the B55 entry (ARPP19) and the B55 hook (P10, ARPP19), (**Figs. 1a-c**). These structures also showed that ARPP19 and FAM122A bind PP2A:B55 via helices rather than extended interactions. The observation that ARPP19 and FAM122A bind B55 almost exclusively via helical interactions suggest that B55-based recruitment is fundamentally distinct from that of its fellow PPP members (while some PP1 regulators had previously shown to bind PP1 using helices in addition to canonical PP1-specific SLiM motifs—Inhibitor-2, spinophilin, NIPP1, MYPT1—it was the SLiM motifs that were essential for PP1 binding). Consistent with this observation, both ARPP19 and FAM122A had shared and distinct interactions with B55 and PP2Ac, highlighting that the interactions between regulators and PP2A:B55 are likely more complex for this holoenzyme.

**Figure 1.**
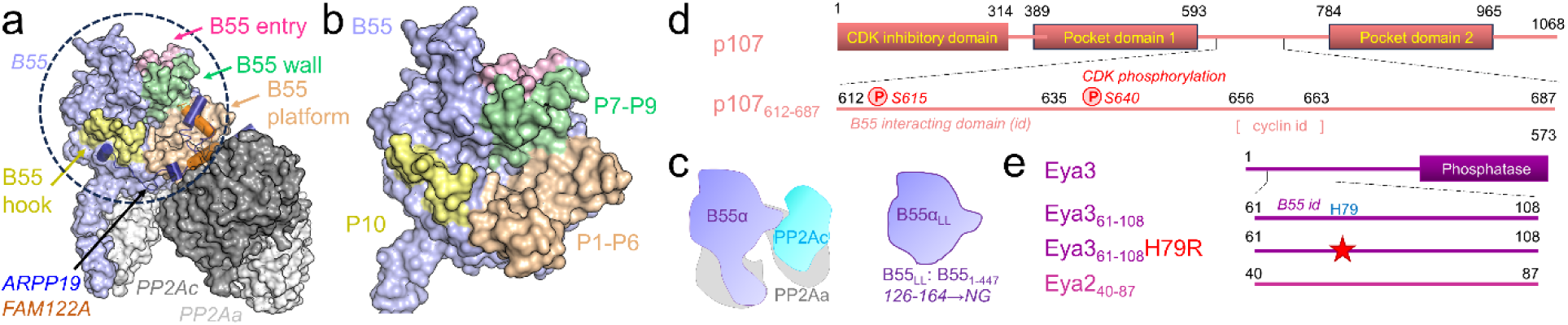
PP2A:B55 and its interactors. **a**. PP2A:B55 (B55, lavender, PP2Aa, light grey and PP2Ac, dark grey) bound to inhibitors ARPP19 (dark blue) or FAM122A (orange). Inhibitors bind B55 via the B55 platform (beige: ARPP19/FAM122A), B55 wall (green: ARPP19/FAM122A), B55 entry (pink: ARPP19) or the B55 hook (yellow; ARPP19). **b**. Closeup of the B55 binding surface, with key interaction pockets (P1-10) indicated. **c**. Cartoon illustrating the PP2A subunits and B55 loopless (B55_LL_); B55_LL_ is monomeric, as it is unable to assemble with PP2Aa or PP2Ac. **d**. Domain structure of p107, with key constructs studied herein indicated. **e**. Domain structure of Eya3 and Eya2 with key constructs studied herein indicated.

Compared to other PPP family members, only limited experimental data is available to fully define the mechanism(s) of PP2A:B55-specific substrate/regulator recruitment. Recently the structure of the PP2A:B55-IER5 substrate complex was described^27^, which showed that while IER5 also uses helices to bind to PP2A:B55, the interaction is very different from that observed for either ARPP19 or FAM122A, increasing the likelihood that there are multiple mechanism(s) used by B55 to recruit substrates/regulators. Here we study the interaction of the PP2A:B55 substrate p107 (**Fig. 1d**) and the scaffold/regulator Eya3 (**Fig. 1e**) to fully define the molecular basis of B55-specific substrate recruitment. p107 is a member of the retinoblastoma family of growth suppressor proteins (RB, p107 & p130) that facilitate the coordinated regulation of cell cycle progression by modulating the E2F family of transcription factors^28,29^. Phosphorylation by CDKs at the intrinsically disordered connector between the two structured pocket domains inactivates p107. PP2A:B55 activates p107 by reversing CDK phosphorylation, a process dependent on the B55-specific recruitment of p107^30,31^. Eya3 is a member of the Eya protein family (Eya1-4) whose members regulate embryonic development via their ability to participate in transcriptional regulation and phosphorylation signaling^32–34^. While it is established that Eya proteins contain a conserved C-terminal tyrosine phosphatase activity, recent work has demonstrated that it also functions as a ser/thr phosphatase via the ability of its N-terminus to bind and recruit PP2A:B55 holoenzymes^35,36^. Recently it was shown that Eya3 recruits PP2A:B55 to c-Myc where it leads to the dephosphorylation of T58 and thus regulates c-Myc stability/activity^37^. Here we use cryogenic electron microscopy (cryo-EM), nuclear magnetic resonance (NMR) spectroscopy, binding and activity assays to molecularly define how PP2A:B55 recruit p107 and Eya3 and understand how these binding interactions facilitate substrate dephosphorylation.

## Results

### Mapping the interaction of p107 and Eya3 with B55 and PP2A:B55

We and others previously showed that p107 binds B55/PP2A:B55 using its intrinsically disordered interdomain region (aa 594-783; with aa 619-625 being essential for B55 binding; pull-down assays with the p107 interdomain region [residues 612-687; hereafter referred to as p107] confirmed PP2A:B55 binding, **Extended Data Figs. 1a,b**)^30^. The 2D [^1^H,^15^N] heteronuclear single quantum coherence (HSQC) spectrum of the B55-interaction p107 interdomain (**Fig. 2a**) confirmed that it is an IDP with modest preferred secondary structure propensities (chemical shift index, CSI) (**Extended Data Fig. 2a**). Similarly, the 2D [^1^H,^15^N] HSQC spectrum of the unbound Eya3 PP2A:B55 interaction domain (aa 62-108), also confirmed that it is an IDP with a single lowly populated α-helix (**Figs. 2b, Extended Data Fig. 2b**).

**Figure 2.**
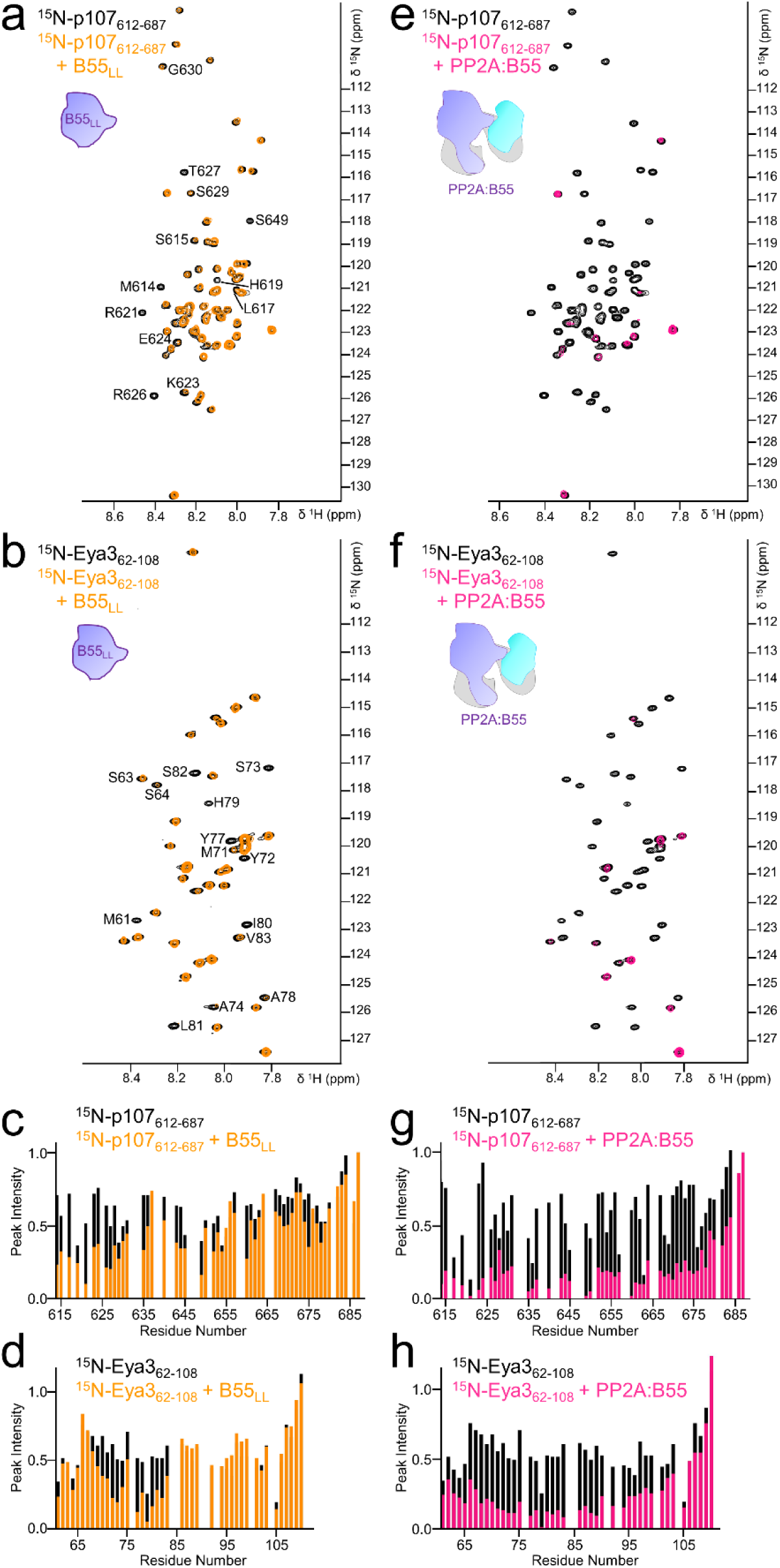
The p107 and Eya3 interactions with B55_LL_ and PP2A:B55 triple complex. **a**. 2D [^1^H,^15^N] HSQC spectrum of ^15^N-labeled p107 with (orange) and without (black) B55_LL_. Peaks missing in the presence of B55_LL_ are labeled. **b**. 2D [^1^H,^15^N] HSQC spectrum of ^15^N-labeled Eya3 with (orange) and without (black) B55_LL_. Peaks missing in the presence of B55_LL_ are labeled. **c**. Plot of peak intensity vs p107 protein sequence for spectra in (a). **d**. Plot of peak intensity vs Eya3 protein sequence for spectra in (b). **e**. 2D [^1^H,^15^N] HSQC spectrum of ^15^N-labeled p107 with (pink) and without (black) PP2A:B55. **f**. 2D [^1^H,^15^N] HSQC spectrum of ^15^N-labeled Eya3 with (pink) and without (black) PP2A:B55. **g**. Plot of peak intensity vs p107 protein sequence for spectra in (e) **h**. Plot of peak intensity vs Eya3 protein sequence for spectra in (f).

Both p107 and Eya3 have been previously shown to bind PP2A:B55 holoenzymes^30,37^. To test this, we incubated ^15^N-labeled (NMR-active) p107 or Eya3 either with or without monomeric B55 (B55_LL_, in which PP2Aa interaction loop, aa 126-164, are replaced with residues ‘NG’ rendering it unable to bind the core PP2A enzyme PP2Aa:PP2Ac) and then overlaid the 2D [^1^H,^15^N] HSQC spectra of p107 or Eya3 to identify peaks (residues) with reduced intensities (peaks with reduced intensities are due to either a direct interaction, a dynamic charge:charge interaction or intermediate timescale conformational exchange; **Figs. 2a-d**). For p107, residues 614-630 experienced the strongest reductions in intensity (**Fig. 2c**). These residues are identical to those previously observed to exhibit reduced intensities with WT B55, confirming not only that p107 binds B55, but also that the B55 PP2Aa binding loop that is missing in B55_LL_, is dispensable for p107 binding. Similar results were obtained with Eya3. Namely, in the presence of B55_LL_, Eya3 residues 68-83 showed reduced intensities (**Fig. 2d**), confirming that Eya3 binds directly to B55_LL_. Notably, the extent of the interaction for both p107 and Eya3 to B55_LL_ is ∼15 residues.

We then repeated the NMR interaction analysis of p107 and Eya3 with the PP2A:B55 holoenzyme (PP2Aa:B55:PP2Ac; purified, active PP2A:B55 is methylated on PP2Ac mL309)^26^. For both interactors, additional residues showed reductions in intensity with PP2A:B55 (**Figs. 2e-h**; p107 aa 614-685; Eya3 aa 63-103), compared to B55_LL_ alone. These data show that residues outside of the B55-interaction domains of p107 and Eya3 interact with either PP2Aa, PP2Ac or both. Furthermore, these data show that both p107 and Eya3 bind directly to the full PP2A:B55 holoenzyme with more than ≥20 residues, suggesting they bind in manners more similar to those observed for ARPP19 and/or FAM122A, rather than the limited interactions observed for substrate/regulators that bind specifically to the B56 subunit in PP2A:B56 holoenzymes using a short B56-specific SLiM (LxxIxE)^7,10^.

### PP2A:B55-p107 and PP2A:B55-Eya3 cryo-EM structures

We determined the cryo-EM structures of PP2A:B55-p107 and PP2A:B55-Eya3 at average resolutions of 2.6 and 2.7 Å, respectively (**Figs. 3a,b, Extended Data Figs. 3-5 and Table 1**). The previously solved cryo-EM structure of PP2A:B55 bound to FAM122A^26^ was used to model the PP2Aa, B55 and PP2Ac subunits. Both complexes adopted the same contracted horseshoe-shaped conformation of PP2Aa observed in the inhibitor bound structures of PP2A:B55-ARPP19 and PP2A:B55-FAM122A. Clear density was observed for the methylated PP2Ac C-terminus (aa 294-309), which, as observed previously, extends across the PP2Aa central cavity to bind an extended pocket at the B55-PP2Aa interface, positioning mL309_C_ to bind a hydrophobic pocket in PP2Aa. These structures confirm that the contracted conformation of PP2A:B55 is largely unaffected by substrate/regulator/inhibitor binding and, instead, is dictated by the methylation state of the C-terminal residues of PP2Ac. In both complexes, unaccounted for density was observed at the B55 platform (**Figs. 3c,d**; the B55 platform is comprised of loops aa 177-179, aa 197-199 and aa 222-231 and contains hydrophobic pockets 1-3 and acidic binding pocket 4; the B55 platform wall is comprised of loops aa 278-287 and aa 336-345 and contains hydrophobic pockets 3 and 4). This density was attributed to p107 (residues 615-635) or Eya3 (residues 72-84), respectively. Although they bind the same surface of B55, they do so via distinct conformations, with p107 forming a helix and Eya3 binding in an extended conformation (**Fig. 3e**).

**Figure 3.**
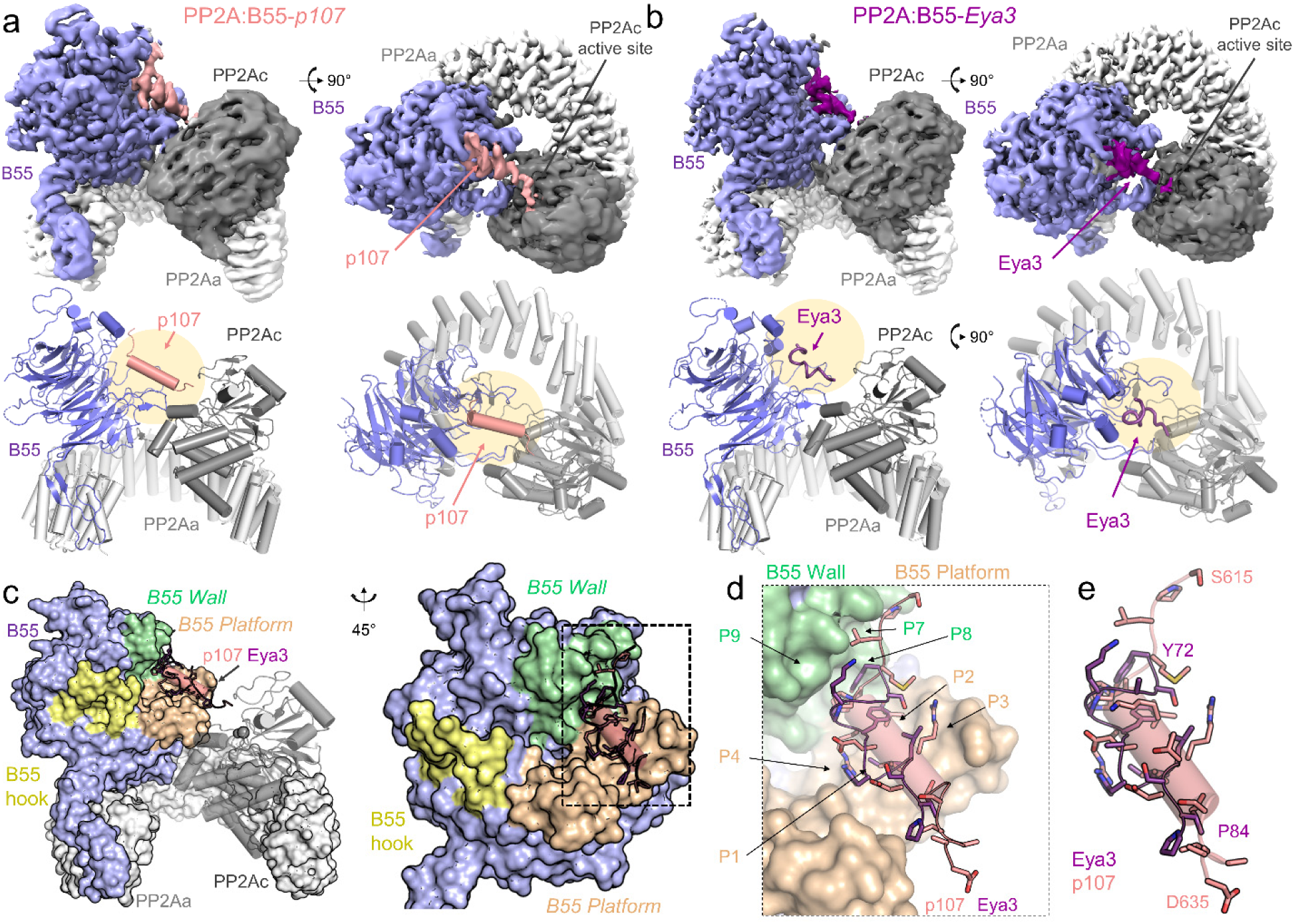
Structures of the PP2A:B55-p107 and PP2A:B55-Eya3 complexes. **a.** Cryo-EM map and model of PP2A:B55-p107. Two views of the map (top) are shown with the corresponding view of the molecular model (bottom). **b**. Cryo-EM map and model of PP2A:B55-Eya3. **c**. Overlay of p107 (salmon) and Eya3 (dark magenta) bound to PP2A:B55 (PP2A colored as in 1a). **d**. Close-up of p107 and Eya3 bound to B55, with interacting residues shown as sticks. Key interacting pockets on B55 are labeled and colored (pockets in the platform, beige; pockets in the wall, green). **e**. Same as (d), but only p107 and Eya3 are shown with the N- and C-terminal residues labeled.

**Table 1:** Cryo-EM data collection, refinement and validation statistics.

|  | **PP2A:B55-p107**<br>**(EMD-45243)(PDB 9C6B)** | **PP2A:B55-Eya3**<br>**(EMD-45292)(PDB 9C7T)** |
| --- | --- | --- |
| **Data collection, processing** |  |  |
| Magnification | 105,000x | 165,000x |
| Voltage (keV) | 300 | 300 |
| Exposure dose | 39.6 | 51.04 |
| Defocus range ($\mu\text{m}$) | -0.55 to -2.6 | -0.6 to -2.0 |
| Pixel size ($\text{\AA}$) | 0.827 | 0.73 |
| Symmetry imposed | $C_1$ | $C_1$ |
| Initial particle images (no.) | 2,800,783 | 874,057 |
| Final particle images (no.) | 195,950 | 107,919 |
| Map resolution ($\text{\AA}$) | 2.6 | 2.7 |
| FSC threshold | 0.143 | 0.143 |
| Map resolution range ($\text{\AA}$) | 1.8 – 24.9 | 1.7 – 18.8 |
| **Refinement** |  |  |
| Initial model used (PDB code) | 8SO0 | 8SO0 |
| Model Composition |  |  |
| Chains | 4 | 4 |
| Atoms | 10736 | 10659 |
| Residues | 1344 | 1333 |
| Ligands | ZN: 1; FE: 1 | ZN: 1; FE: 1 |
| R.m.s. deviations |  |  |
| Bond lengths ($\text{\AA}$) | 0.003 | 0.004 |
| Bond angles ($^\circ$) | 0.475 | 0.538 |
| B factors ($\text{\AA}^2$) min/max/mean | | |
| Protein | 85.0/190.05/128.9 | 43.4/140.4/81.2 |
| Ligand | 160.4/164.9/162.7 | 112.3/113.2/112.7 |
| Map-model fit ($\text{\AA}$) | 2.8 | 2.7 |
| FSC | 0.5 | 0.5 |
| **Validation** |  |  |
| MolProbity score | 1.26 | 1.32 |
| Clashscore | 3.89 | 4.01 |
| Rotamer outliers (%) | 0.00 | 0.00 |
| Ramachandran plot (%) |  |  |
| Outliers | 0.00 | 0.08 |
| Allowed | 2.4 | 2.7 |
| Favored | 97.6 | 97.3 |
| Rotamer outliers (%) | 0.00 | 0.00 |
| Cβ outliers (%) | 0.00 | 0.00 |

### PP2A:B55 recruitment of p107

p107 binds B55 as a helix, with the residues N-terminal to the helix binding the B55 wall and those C-terminal extending towards the PP2Ac catalytic site (**Fig. 4a**). The interaction of the p107 N-terminus (^615^SPLMHP^620^) with the B55 wall is anchored by L617_p107_, which binds B55 hydrophobic pocket P7 (F280_B55_, F281_B55_, I284_B55_, Y337_B55_, F343_B55_; **Fig. 4b**) and H619_p107_, which binds B55 hydrophobic/polar pocket P8 (S247_B55_, E283_B55_, I284_B55_, S287_B55_, F343_B55_, K345_B55_; **Fig. 4c**). At P620_p107_, p107 kinks to bind across the base of the B55 platform, forming a 3-turn helix (621-632) that extends towards the PP2Ac catalytic pocket, with p107 residues 633-635 abutting PP2Ac (**Fig. 3a**). While residues 634-687 of p107 were not sufficiently ordered to be modeled, our NMR data suggest that they contribute to binding via a dynamic (fuzzy) charge:charge interaction^5,23,38^ (**Fig. 2g**).

**Figure 4.**
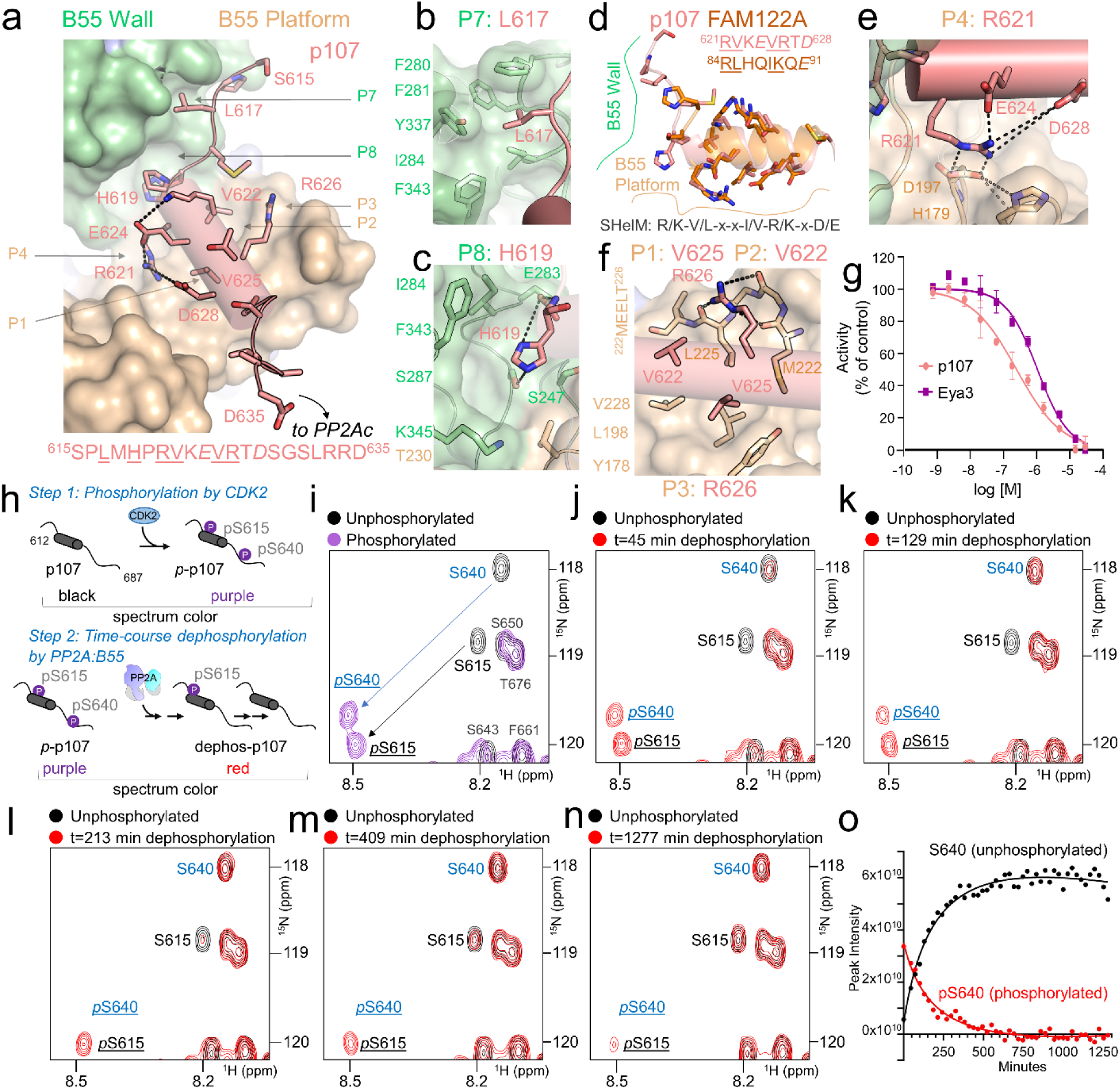
PP2A:B55 preferentially dephosphorylates p107 CDK2/Cyclin A2 phosphosite *p*S640. **a** PP2A:B55-p107 showing that p107 (salmon, shown as sticks) binds the B55 platform (beige) and wall (green). B55 interaction pockets occupied by p107 are labeled and their positions indicated by arrows. Key p107 interacting residues are labeled. The interaction sequence is shown below, with residues that mediate intermolecular contacts underlined and those important for intramolecular contacts in italics. **b**. p107 L617 (salmon) binds the B55 P7 hydrophobic pocket (P5 pocket residues shown as sticks). **c**. p107 H619 (salmon) binds the B55 P8 hydrophobic/polar pocket (P6 pocket residues shown as sticks; polar interactions shown with a black dotted line). **d**. Overlay of p107 (salmon) and the FAM122A B55 binding helix (orange), with side chains shown as sticks. Structural alignment of p107 and FAM122A interacting residues are also shown. **e**. p107 R621 (salmon) binds the B55 P4 acidic pocket (P4 pocket and key p107 residues shown as sticks; the multiple inter- and intramolecular interactions that stabilize the conformation are shown as black dotted lines). **f**. p107 V622 and V625 bind the B55 hydrophobic P2 and P1 pockets (P1/P2 pocket residues shown as sticks), respectively, while p107 R626 binds the hydrophobic/polar P3 pocket (P3 pocket residues shown as sticks with hydrogen bonds between R626 and the ^222^MEELT^226^ carbonyls indicated by dashed lines). **g**. PP2A:B55 inhibition by p107 and Eya3 (mean ± s.d.; n = 3 experimental replicates). IC_50_ values are reported in Extended Data Table 1. **h**. Cartoon illustrating p107 phosphorylation (CDK2/Cyclin A2) and subsequent time-course of dephosphorylation (PP2A:B55) assays. **i**. 2D [^1^H,^15^N] HSQC spectrum of ^15^N-labeled unphosphorylated p107 (black) overlaid on CDK2/Cyclin A2 phosphorylated ^15^N-labeled p107 (purple); S615 and S640 are phosphorylated, with shifted peaks labeled. **j-n**. 2D [^1^H,^15^N] HSQC spectrum of unphosphorylated ^15^N-labeled p107 (black) overlaid with that of CDK2/Cyclin A2 phosphorylated ^15^N-labeled p107 incubated with PP2A:B55 (red) for the following timepoints in minutes: 45 (j), 129 (k), 213 (l), 409 (m) and 1277 (n). **o**. Changes in peak intensities of ^15^N-labeled p107 residues *p*S640 (red) and S640 (black).

Previously, we showed that the PP2A:B55-specific inhibitor FAM122A binds B55 using a helix and that the FAM122A residues that mediate B55-binding (using a short helical motif, or SHelM, defined as R-L/V-x-x-I/V-K/R-x-E/D)^26^ are conserved in B55-binding domain of p107, suggesting they bind using similar mechanisms. The structure of p107 bound to PP2A:B55 confirms that this is the case, with p107 ^621^<u>RV</u>KE<u>VR</u>T<u>D</u>^628^ residues binding the B55 platform in a manner identical to FAM122A ^84^<u>RL</u>HQ<u>IK</u>Q<u>E</u>^91^ (**Fig. 4d**). Like R84_FAM122A_, R621_p107_ forms a bidentate salt bridge with D197_B55_ (pocket P4), a residue previously shown is essential for p107 binding^30^. The conformation of R621_p107_ is stabilized by intramolecular electrostatic interactions with E624_p107_ and D628_p107_ and an intermolecular π-stacking interaction with H179_B55_ (pocket P4) (**Fig. 4e**). Similarly, V622_p107_ and V625_p107_ anchor p107 to the B55 platform via the same hydrophobic pockets used by L85_FAM122A_ and I88_FAM122A_ (bound to pockets P2 and P1, respectively: P2: L225_B55_, V228_B55_; P1 Y178_B55_, L198_B55_, M222_B55,_ L225_B55_, V228_B55_). Finally, R626_p107_, like K89_FAM122A_, forms multiple hydrogen bonds with the carbonyls of B55 loop ^222^MEELT^226^ (P3) (**Fig. 4f**). C-terminal to the B55 binding helix, p107 residues 631-633 extend with less well-defined density towards the PP2Ac active site, suggesting p107 may transiently hinder active site access. To test if p107 binding alters PP2Ac catalytic activity, we performed IC_50_ measurements using DiFMUP as a substrate. The data show that, as predicted, p107 inhibits DiFMUP dephosphorylation (IC_50_: 245 ± 27 nM; **Extended Data Table 1; Fig. 4g**).

### p107 dephosphorylation by PP2A:B55

While p107 slows the dephosphorylation of the active site phosphomimetic DiFMUP pseudo-substrate, p107 itself is a protein substrate of PP2A:B55. p107 is phosphorylated by multiple kinases, including the proline-directed ser/thr cyclin-dependent kinase 2 (CDK2)/Cyclin A2. Of the 14 S/T residues in the p107 interdomain (aa 612-687), S615_p107_, S640_p107_ and S650_p107_ are followed by a proline, and all have been observed to be phosphorylated in multiple phosphoproteomics studies (PhosphoSitePlus)^39^. To identify the p107 residues phosphorylated by CDK2/Cyclin A2, we incubated ^15^N-labeled p107 with purified CDK2/Cyclin A2 and then identified the phosphorylated residues using NMR spectroscopy (**Fig. 4h**, *step 1*). The 2D [^1^H,^15^N] HSQC spectrum of *phospho-*p107 (*p*p107) showed that CDK2/Cyclin A2 fully phosphorylates ^615^SP^616^ and ^640^SP^641^ (**Fig. 4i**). In contrast, ^650^SP^651^ was not phosphorylated, likely due to its proximity to p107’s Cyclin A2 recruiting motif (^654^GSAKRRLFGE^663^)^40^. We then monitored the kinetics of *p*S615_p107_ and *p*S640_p107_ dephosphorylation by PP2A:B55 by following the intensity changes in the phosphorylated (*p*S615_p107_ and *p*S640_p107_) or unphosphorylated (S615_p107_ and S640_p107_) HN/N cross peaks in the 2D [^1^H,^15^N] HSQC spectrum (**Figs. 4h**, *step 2*, **4j-4o**). While *p*S640_p107_ is rapidly dephosphorylated (**Figs. 4j-l, 4o**), the dephosphorylation of *p*S615_p107_ is comparatively slow, with little to no dephosphorylation observed until *p*S640_p107_ is almost completely dephosphorylated (**Figs. 4l-n, Extended Data Figs 6 and 7**). These results are fully consistent with the cryo-EM structure of the PP2A:B55-p107 substrate complex. Namely, S615_p107_ binds directly to B55 in a pocket more than 40 Å away from the PP2Ac active site making it unable to be dephosphorylated when bound to B55. In contrast, S640_p107_ is positioned next to the PP2Ac active site when bound to B55, making it ideally located for rapid dephosphorylation. Together, these data suggest that pS640_p107_ is the primary phosphosite targeted by PP2A:B55.

### PP2A:B55 recruitment of Eya3

While p107 bound B55 as predicted, the lack of the B55-specific SHelM in Eya3, coupled with the lack of a preferred secondary structure in the unbound state (**Extended Data Fig. 2b**), suggested that it may bind B55 via a distinct mechanism. The structure of PP2A:B55-Eya3 confirms that Eya3 binds B55 in an extended, rather than helical, manner (**Fig. 5a**). Despite this difference in conformation, the B55 interaction pockets engaged by Eya3 mirror those used by other B55-specific regulators. First, Y72_Eya3_, K75_Eya3_ and Y77_Eya3_ bind hydrophobic pockets P8, P9 and P2, respectively (**Fig. 5b**), anchoring the N-terminus of Eya3 to the center of B55. This positions Eya3 H79_Eya3_ (a residue that when mutated to Ala has been previously shown to abolish B55 binding^37^) to bind the B55 acidic pocket, P4. H79_Eya3_ binds D197_B55_ from the opposite direction compared to R621_p107_ in the p107 complex, allowing it to form a nearly perfect π-stacking interaction with H179_B55_ (**Fig. 5c**). This interaction is further stabilized by I80_Eya3_ (bound to P1) and L81_Eya3_ (bound to P3) (**Fig. 5c**). Finally, as observed for p107, the C-terminal Eya3 residues ^82^SVP^84^ extend towards the PP2Ac active site. To test if Eya3 influences the PP2Ac catalytic activity against a small phosphomimic, we again performed IC_50_ measurements using DiFMUP as a substrate, which showed weak inhibition (IC_50_: 1102 ± 84 nM; **Extended Data Table 1; Fig. 4g**).

**Figure 5.**
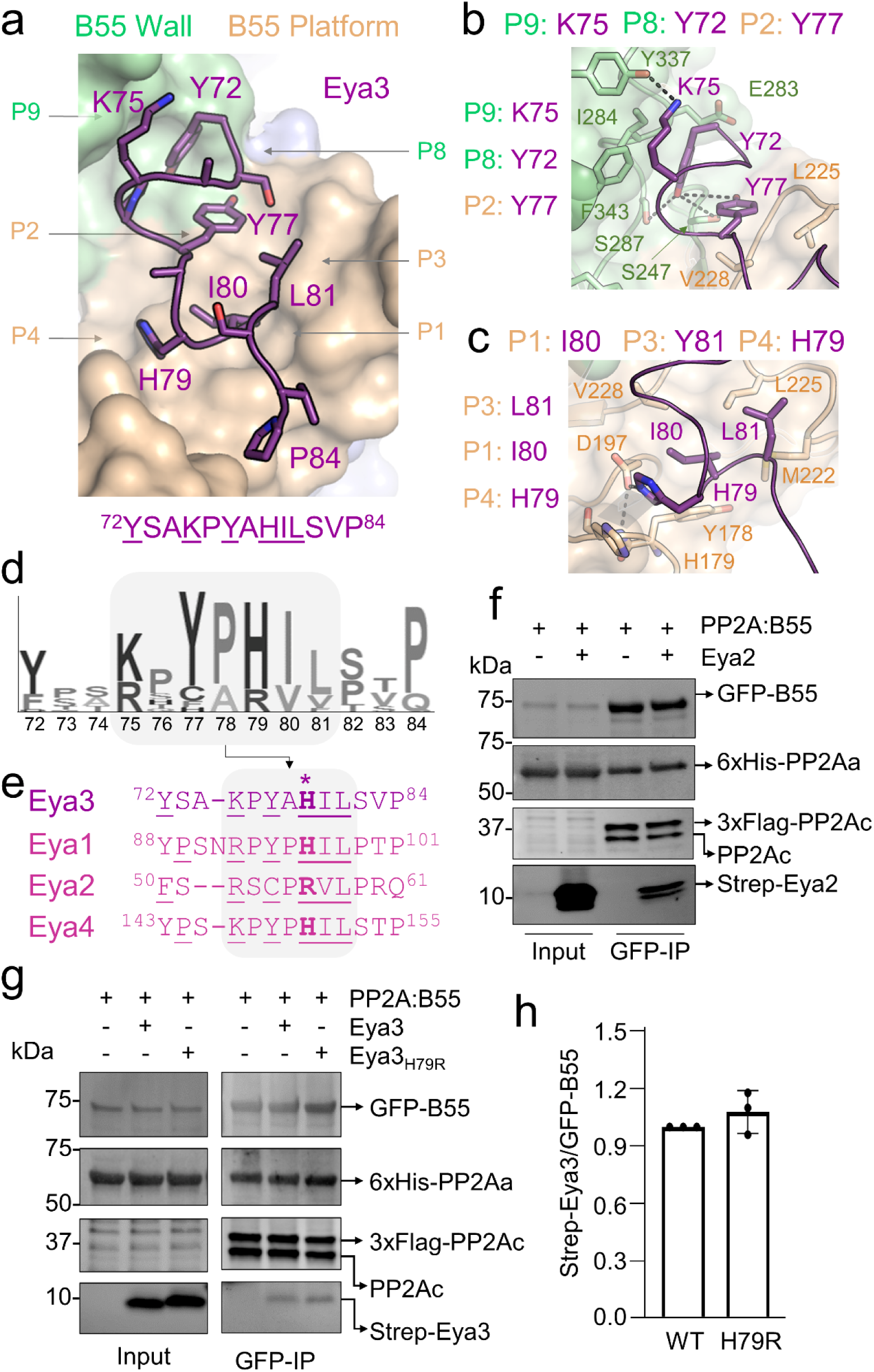
Eya3 binds B55 in an extended conformation via a conserved interaction sequence. **a.** PP2A:B55-Eya3 showing that Eya3 (purple, shown as sticks) binds the B55 platform (beige) and wall (green). B55 interaction pockets occupied by Eya3 are labeled and their positions indicated by arrows. Key Eya3 interacting residues are labeled. The interaction sequence is shown below, with residues that mediate intermolecular contacts underlined. **b**. Interactions between Eya3 (purple) and B55 (wall, green; platform; beige) at pockets P2, P8 and P9 shown. Interacting residues shown as sticks and labeled. Hydrogen bond/salt bridge interactions shown as black dotted lines. **c**. Interactions between Eya3 (purple) and B55 (platform; beige) at pockets P1, P3 and P4 shown. Interacting residues shown as sticks and labeled. Hydrogen bond/salt bridge interactions shown as black dotted lines. The π-stacking interaction between H79_Eya3_ and H179_B55_ indicated by grey shading. **d**. Sequence conservation logo for the Eya3 B55-interaction domain identified using Jackhmmer^54^. The key interaction region is highlighted in grey. **e**. Sequence alignment of the Eya3 B55-interaction domain with the corresponding residues in Eya1, Eya2 and Eya4. Residues corresponding to H79_Eya3_ are in bold. **f**. Expi293F cell lysates expressing GFP-B55, 3xFLAG-PP2Ac and 6xHis-PP2Aa were incubated with or without purified Eya2 (residues 40-84; includes the putative B55 interaction domain), immunopurified using GFP-trap beads (EGFP nanobodies coupled to agarose resin) and isolated proteins detected by immunoblot. Results representative of 3 independent experiments. See also Extended Data Fig. 1a. **g**. Same as f, except the lysate was incubated with either purified Eya3 or Eya3_H79R_. (residues 62-108). **h**. Quantification (mean ± s.d. n = 3 experimental replicates) of Eya3 binding observed in (g), with individual data points shown as dots.

The Eya3 residues that mediate the core interaction with B55 are also highly conserved, especially Y77_Eya3_ and H79_Eya3_ (**Fig. 5d**). In particular, the latter residue was previously shown to be essential for PP2A:B55 binding, as the Eya3 H79A variants severely reduces its ability to recruit PP2A:B55 and, in turn, nearly abolishes its Thr-specific phosphatase activity^36,37^. Comparing the Eya3 B55-interacting sequence with the remaining Eya isoforms (Eya1, Eya2 and Eya4) showed that this region is conserved among the Eya family, with the greatest differences in sequence observed for Eya2, the isoform previously identified to exhibit the highest Thr phosphatase activity (**Fig. 5e**)^37^. We first tested if the Eya2 N-terminal IDP (aa 40-84), which includes the putative B55 interaction residues ^50^<u>F</u>S<u>R</u>S<u>C</u>P<u>RVL</u>PRQ^61^, binds directly to PP2A:B55 using pull-down assays. The data shows that Eya2 residues 40-84 are sufficient to bind and recruit PP2A:B55 (**Fig. 5f**). We then generated the Eya3 H79R variant, in which H79_Eya3_ was replaced with equivalent residue in Eya2 (R56_Eya2_), to test if an arginine impacts PP2A:B55 recruitment. The data show that the B55 binding is statistically equivalent (**Figs. 5g, h**) and thus that either residue bind the B55 P4 binding pocket effectively. Consistent with this observation, modeling an arginine in this position in the PP2A:B55-Eya3 structure shows that H79R is, like H79, optimally positioned to form both a π-stacking interaction with H179_B55_ and a salt bridge with D197_B55_.

## Discussion

A molecular understanding of how PP2A:B55 recruits its substrates and regulators is necessary to identify its endogenous substrates on a proteome-wide scale. The cryo-EM structures of PP2A:B55 in complex with two B55-specific protein inhibitors, FAM122A and ARPP19, showed that they interact with PP2A:B55 in unexpected manners^26^. First, while most PPP-specific regulators and substrates bind their cognate PPPs in extended conformations via SLiMs, FAM122A and ARPP19 bind PP2A:B55 using helices (SHelMs). Second, the structures also revealed that while FAM122A and ARPP19 bind extensively to both B55 and PP2Ac, they do so via very distinct conformations. In addition, the recently determined structure of the PP2A:B55-IER5 complex^27^ showed that IER5 forms a helix-turn-helix bundle that engages B55 via a third conformation. Together, these data suggest that regulators and substrates likely use diverse strategies to bind and recruit PP2A:B55. To define the mechanisms used by substrates and regulators to recruit PP2A:B55 and to understand how these interactions direct PP2A:B55 activity, we determined the cryo-EM structures of a B55-specific substrate, p107, and a regulator, Eya3, bound to PP2A:B55 and determined how these interactions influence substrate dephosphorylation.

Our data showed that both p107 and Eya3 bind to B55 using B55 interaction surfaces also used by FAM122A, ARPP19 and IER5; that is, the B55 wall and B55 platform (**Fig. 6a**). Thus, while two interaction surfaces of B55 engage multiple regulators (wall, platform), others (thus far) are used exclusively by ARPP19 (entry, hook); future work will be needed to determine if these latter interaction surfaces are truly specific to ARPP19 or, alternatively, are used by other regulators and/or substrates. In addition, only the inhibitors FAM122A and ARPP19 also stably bind the PP2Ac active site (**Fig. 6b**). Notably, while p107 and Eya3 bind the same interaction surfaces, they do so via distinct conformations. p107 adopts a helical conformation, overlapping almost perfectly with the B55 interaction helix of FAM122A (this was predicted as their sequences define the only known B55-specific SHelM, R-L/V-x-x-I/V-K/R-x-E/D)^26^. In contrast, Eya3 binds B55 in an extended manner. Superimposing all five B55-regulator/substrate complexes shows that only p107 and FAM122A bind B55 via identical conformations; the rest of the regulators bind the wall and platform pockets via distinct conformations (**Figs. 6c-g**). This suggests that B55 interaction proteins (substrates/regulators) likely use different combinatorial strategies to recruit B55 via the wall and platform binding pockets, making predicting novel B55-substrates and regulators from sequence alone challenging.

**Figure 6.**
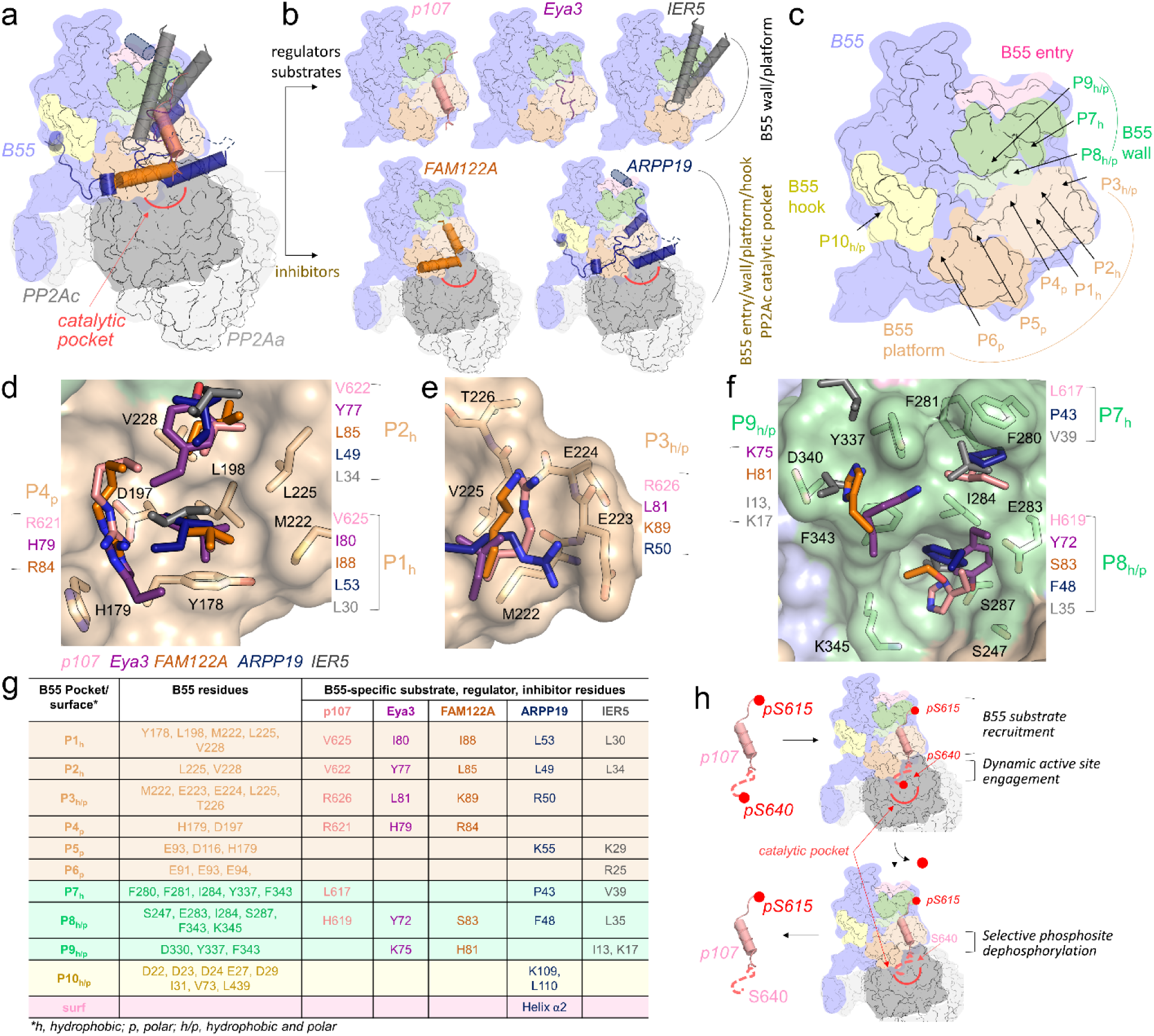
Combinatorial mechanisms used by B55-specific regulators, substrates and inhibitors contribute to PP2A:B55 phosphosite selectivity. **a**. Overlay of PP2A:B55 (B55 in lavender with interaction sites colored [platform, beige; wall, green; entry, pink; hook, yellow]; PP2Ac, dark grey; PP2Aa, white) bound to p107 (salmon), eya3 (purple), FAM122A (orange), ARPP19 (dark blue) and IER5 (gray). The location of the PP2Ac catalytic pocket is indicated in red. **b**. Same as (a), but individual complexes, separated by regulators/substrates and inhibitors. **c**. Location of individual binding pockets on B55, colored as in (a). **d**. B55 interactor residues bound at B55 pockets P1, P2 and P4. B55 pocket residues, beige sticks; B55 interactor residues, sticks and colored as in (a). B55 interactor residues labeled by color as in (a); B55 pocket residues labeled in black. **e**. Same as (d) but for B55 pocket P3. **f.** Same as (d) but for B55 pockets P7-P9. **g**. B55 pockets, with residues defining the pockets indicated and the corresponding B55 interacting residues listed. **h**. Cartoon illustrating how B55-specific recruitment contributes to PP2A:B55 phosphosite dephosphorylation selectivity.

In addition to defining substrate binding, the PP2A:B55-p107 structure, coupled with atomic resolution dephosphorylation assays, provided key insights into the role of B55-mediated recruitment for directing PP2A:B55 phosphosite specificity. We showed that while the p107 interdomain is robustly phosphorylated by CDK2/Cyclin A2 on two residues, S615_p107_ and S640_p107_, it is only pS640_p107,_ and not pS615_p107_, that is readily dephosphorylated by PP2A:B55 (**Fig. 4**). This preference in phosphosite dephosphorylation is readily explained by the PP2A:B55-p107 structure. Namely, in the bound state, S615_p107_ is associated with B55 at the wall interaction surface, more than 40 Å away from the PP2Ac catalytic pocket. In contrast, in the bound state, S640_p107_ is positioned directly adjacent to the active site (**Fig. 6h, upper panels**), making substrate engagement dephosphorylation readily achievable. In this way, B55-binding ensures that only specific phosphosites (i.e., pS640_p107_) are preferentially dephosphorylated (**Fig. 6h, lower panels**).

Finally, our work also highlights the importance of combining cryo-EM with NMR spectroscopy to have a comprehensive understanding of how IDP regulators and substrates engage their cognate folded binding partners. By studying the ensemble of the IDP regulators in their unbound and bound forms, we showed that while FAM122A and ARPP19 have a preference for prepopulated secondary structures in their unbound forms, this behavior was not observed (thus far) for regulators or substrates. The data also showed while cryo-EM identified the most highly populated conformations of substrates and regulators in the bound state, residues outside this region are also critical for binding (**Figs. 2e, f**)^41^. Indeed, for both p107 and Eya3, many residues beyond the core B55 interaction domains showed a loss of intensity in the presence of PP2A:B55, likely corresponding to a dynamic, charge-charge (fuzzy) interaction with the surface of PP2Ac or PP2Aa^42–44^. These data are fully consistent with previous PP2A:B55 work. For example, in addition to the well-ordered interactions of ARPP19 with the B55 wall, platform and hook, cryo-EM and especially NMR spectroscopy showed that a pre-populated helix in the free state also dynamically engages the B55 entry interaction surface, an interaction that is essential for the simultaneous binding of ARPP19 and FAM122A to PP2A-B55^26^. The importance of dynamic interactions in regulator/substrate recruitment by PPPs is true not only for PP2A:B55, but also PP2A:B56,^23^ calcineurin^5^ and PP1^38^. NMR spectroscopy is the only technique that captures these dynamic interactions at atomic resolution; thus, combining NMR spectroscopy with other high-resolution methods is essential to obtain a comprehensive understanding of how PPP-specific regulators and substrates recruit their cognate PPP.

Together, these studies provide a molecular understanding of regulator and substrate recruitment of the PP2A:B55 holoenzyme. Because of the key regulatory functions of PP2A:B55 in mitosis and DNA damage repair, these data provide a roadmap for characterizing disease-associated mutations and pursuing new avenues to therapeutically target this complex, by individually blocking a subset of regulators that use different B55 interaction sites.

## Methods

### Bacterial protein expression

PP2Aa_9-589_, p107_612-687_, EYA2_40-84EEE_ and Eya3_62-108EE_ were subcloned into pTHMT containing an N-terminal His_6_-tag followed by maltose binding protein (MBP) and a tobacco etch virus (TEV) protease cleavage site. For expression, plasmid DNAs were transformed into *E. coli* BL21 (DE3) cells (Agilent). Freshly transformed cells were grown at 37°C in LB broth containing kanamycin antibiotics (50 µg/ml) until they reached an optical density (OD_600_) of ∼0.8. Protein expression was induced by addition of 1 mM β-D-thiogalactopyranoside (IPTG) to the culture medium, and cultures were allowed to grow overnight (18-20 hours, 250 rpm shaking) at 18°C. Cells were harvested by centrifugation (8000 *x*g, 15 min, 4°C) and stored at - 80°C until purification. Expression of uniformly ^13^C- and/or ^15^N-labeled protein was carried out by growing freshly transformed cells in M9 minimal media containing 4 g/L [^13^C]-D-glucose and/or 1 g/L ^15^NH_4_Cl (Cambridge Isotopes Laboratories) as the sole carbon and nitrogen sources, respectively. The Eya3 H79R variant was generated by site-directed mutagenesis, sequence verified and expressed as described above. For pull-down experiments p107, EYA2, EYA3 and EYA3_H79R_ were expressed with a C-terminal StrepII tag for detection using a Strep antibody (Genscript, A01732, 1:1000).

### Cell culture

Expi293F cells were obtained from ThermoFisher (cat# A14527) and grown in HEK293 Cell Complete Medium (SMM 293-TII, Sino Biological cat# M293TII). For transient overexpression of B55 and PP2Ac constructs, cells were transfected using polyethyleneimine (PEI) transfection reagent. For Western blot and immunoprecipitation studies, whole cell extracts were prepared by lysing cells in ice-cold lysis buffer (20 mM Tris pH 8.0, 500 mM NaCl, 0.5 mM TCEP, 1 mM MnCl_2_, 0.1% Triton-X100, Phosphatase inhibitor cocktail [ThermoFisher]), sonicating and clearing the lysate by centrifuging at 15,000 *x*g for 20 min at 4°C. Total protein concentrations were measured using the Pierce 660 Protein Assay Reagent (ThermoFisher).

### Mammalian protein expression

Full length B55_1-477_ was cloned into pcDNA3.4 including an N-terminal green fluorescence protein (GFP) followed by a TEV cleavage sequence. Full length PP2Ac_1-309_ was cloned into pcDNA3.4 with an N-terminal Strep tag followed by a TEV cleavage sequence. B55 loop-less (B55_LL_), in which B55 residues 126-164 that interact directly with PP2Aa, were removed, and replaced with a single NG linker (**Fig. 1c**), was cloned into pcDNA3.4 with an N-terminal green fluorescence protein (GFP) followed by a TEV cleavage sequence. All plasmids were amplified and purified using the NucleoBond Xtra Maxi Plus EF (Macherey-Nagel). B55_WT_ and B55_LL_ were individually expressed in Expi293F cells (ThermoFisher). B55_1-477_ and PP2Ac_1-309_ were co-expressed in Expi293F cells at a 1:2 DNA ratio.

Transfections were performed in 500 mL medium (SMM293-TII, Sino Biological) in 2 L flasks using polyethylenimine (PEI, Polysciences) reagent according to the manufacturer’s protocol in an incubator at 37°C and 8% CO_2_ under shaking (125 rpm). On the day of transfection, the cell density was adjusted to 2.8 X10^6^ cells/mL using fresh SMM293-TII expression medium. DNA of PP2Ac and B55 (2:1 ratio) were diluted in Opti-MEM Reduced Serum Medium (ThermoFisher). Similarly, in a separate tube, PEI 3x the amount of DNA was diluted in the same volume of Opti-MEM Reduced Serum Medium (ThermoFisher). The DNA and PEI mixtures were combined and incubated for 10 min at room temperature, before being added to the cell culture. Valproic acid (2.2 mM final concentration, Sigma) was added to the cells 4 h after transfection and 24 h after transfection sterile-filtered glucose (4.5 mL per 500 mL cell culture, 45%, glucose stock) was added to the cell culture flasks to boost protein production. Cells were harvested 48 hrs after transfection by centrifugation (2,000 x*g* for 20 minutes, 4°C) and stored at −80°C.

### p107, Eya3 and Eya2 purification

Cell pellets were resuspended in ice-cold lysis buffer (50 mM Tris pH 8.0, 500 mM NaCl, 5 mM Imidazole, 0.1% Triton X-100, protease inhibitor tablet [ThermoFisher]), lysed by high-pressure cell homogenization (Avestin Emulsiflex C3). Cell debris was pelleted by centrifugation (42,000 x*g*, 45 min, 4°C), and the supernatant was filtered with 0.22 µm syringe filters (Millipore). The proteins were loaded onto a HisTrap HP column (Cytiva) pre-equilibrated with Buffer A (50 mM Tris pH 8.0, 500 mM NaCl, 5 mM imidazole) and eluted using a linear gradient (0-60%) with Buffer B (50 mM Tris pH 8.0, 500 mM NaCl and 500 mM imidazole). Fractions containing the protein were pooled and dialyzed overnight at 4°C with TEV protease (in-house; his_6_-tagged) to cleave the His_6_-MBP tag. Following cleavage, the sample was loaded under gravity onto Ni^2+^-NTA beads (Prometheus) pre-equilibrated with Buffer A, the flow through and wash A fractions were collected, and then heat purified by incubating the samples at 80°C for 20 min Samples were centrifuged at 15,000 *x*g for 10 min to remove precipitated protein. Supernatant was concentrated and purified using size exclusion chromatography (SEC; Superdex 75 16/60 [Cytiva]) in either p107 NMR buffer (10 mM Na/PO_4_ pH 6.3, 150 mM NaCl, 0.5 mM TCEP), Eya3 NMR buffer (10mM Na/PO_4_ pH 6.5, 200 mM NaCl, 0.5 mM TCEP) or IC_50_ assay buffer (20 mM Tris pH 8.0, 150 mM NaCl, 0.5 mM TCEP). Purified samples were again heat purified (80°C for 5 minutes), and were either directly used for NMR data collection or flash frozen and stored at −80°C.

### PP2Aa purification

Cell pellets expressing PP2Aa_9-589_ were resuspended in ice-cold lysis buffer (50 mM Tris pH 8.0, 500 mM NaCl, 5 mM Imidazole, 0.1% Triton X-100, EDTA-free protease inhibitor tablet [ThermoFisher]), lysed by high-pressure cell homogenization (Avestin Emulsiflex C3). Cell debris was pelleted by centrifugation (42,000 x*g*, 45 min, 4°C), and the supernatant was filtered with 0.22 µm syringe filters (Millipore). The proteins were loaded onto a HisTrap HP column (Cytiva) pre-equilibrated with Buffer A (50 mM Tris pH 8.0, 500 mM NaCl, 5 mM imidazole) and eluted using a linear gradient (0 to 40%) with Buffer B (50 mM Tris pH 8.0, 500 mM NaCl and 500 mM imidazole). Fractions containing the protein were pooled and dialyzed overnight at 4°C with TEV protease (in-house; his_6_-tagged) to cleave the His_6_-MBP-tag and loaded under gravity onto Ni^2+^-NTA beads (Prometheus) pre-equilibrated with Buffer A. Flow through and wash A fractions were collected, concentrated and loaded onto QTrap HP column (Cytiva) for further purification. The proteins were eluted with a 100 mM – 1 M salt gradient (Buffer A: 20 mM Tris pH 8.0, 100 mM NaCl, 0.5 mM TCEP; Buffer B: 20 mM Tris pH 8.0, 1 M NaCl, 0.5 mM TCEP). PP2Aa fractions were concentrated and further purified using size exclusion chromatography (SEC; Superdex 200 26/60 [Cytiva]) in Assay buffer (20 mM Tris pH 8.0, 150 mM NaCl, 0.5 mM TCEP). Samples were either directly used or flash frozen and stored at −80°C.

### CDK2/Cyclin A2 kinase expression and purification

CDK2 was subcloned into pGEX3C. Cyclin A2_156-429_ was subcloned into pET16b. The plasmids were transformed separately into *E. coli* BL21 (DE3) cells (Agilent). Freshly transformed cells were grown at 37°C in LB broth containing carbenicillin (50 µg/ml) until they reached an optical density (OD_600_) of ∼0.8. Protein expression was induced by addition of 1 mM IPTG to the culture medium, and cultures were allowed to grow overnight (18-20 hours, 250 rpm shaking) at 18°C. Cells were harvested by centrifugation (8000 *x*g, 15 min, 4°C) and stored at −80°C until purification. Cell pellets expressing CDK2 and Cyclin A2 were resuspended in lysis buffer (50 mM Tris pH 8.0, 500 mM NaCl, 5 mM Imidazole, 0.1% Triton X-100, protease inhibitor tablet [ThermoFisher]) and combined prior to high-pressure cell homogenization (Avestin Emulsiflex C3). Cell debris was pelleted by centrifugation (42,000 x*g*, 45 min, 4°C), and the supernatant was filtered with 0.22 µm syringe filters (Millipore). The proteins were loaded onto a HisTrap HP column (Cytiva) pre-equilibrated with Buffer A (50 mM Tris pH 8.0, 500 mM NaCl, 5 mM imidazole) and eluted using a linear gradient (0 to 60%) with Buffer B (50 mM Tris pH 8.0, 500 mM NaCl and 500 mM imidazole). Fractions containing both proteins were pooled and dialyzed overnight at 4⁰C in Size Exclusion Chromatography (SEC) Buffer (20 mM Tris pH 7.5, 500 mM NaCl, 1 mM DTT). The complex was concentrated and further purified using SEC (Superdex 200 26/60 [Cytiva]). Fractions containing the proteins were pooled and dialyzed in dialysis buffer (20 mM Tris pH 7.5, 150 mM NaCl, 1 mM MgCl_2_, 0.005% Tween-20) overnight at 4°C. The CDK2/Cyclin A2 complex was mixed with 50% glycerol and stored at −80°C.

### p107 phosphorylation

Purified ^15^N-labeled-p107 (25 μM) was incubated with CDK2/Cyclin A2 kinase (500:1 ratio) in phosphorylation buffer (100 mM Tris pH 7.5, 2 mM DTT, 10 mM MgCl_2_) with 500 µM of ATP (Sigma) for phosphorylation. The kinase reaction was incubated at 37⁰C for 1 hour. Phosphorylated p107 (*p*p107) was immediately heat purified by incubating *p*p107 at 80°C for 10 min. The sample was centrifuged at 15,000 *x*g for 10 min to remove precipitated CDK2/Cyclin A2 and either immediately used for experiments or flash frozen and stored at −80⁰C. Complete phosphorylation was confirmed by monitoring the chemical shift changes of the phosphorylated serine residue(s) using NMR spectroscopy (2D [^1^H,^15^N] HSQC).

### p107, Eya3 and variants and Eya2 interaction with the PP2A:B55 complex

Purified p107 (∼25 μg), Eya3, Eya3_H79R_ or Eya2 (∼25 μg) alone were mixed with Expi293F whole-cell extracts expressing B55, PP2Ac and PP2Aa constructs. Input samples were collected prior to incubation with GFP-Trap agarose beads (**Extended Data** Fig. 1a). GFP-tagged B55 and associated proteins were captured by incubating equal amounts of total protein (∼500 μg) for each condition with GFP-Trap agarose beads (prepared as described in ‘EGFP–nanobody protein expression, purification, and immobilization onto agarose beads’ in Methods) at 4°C for 2 h. Following 3 washes with wash buffer (20 mM Tris pH 8.0, 500 mM NaCl, 0.5 mM TCEP, 1 mM MnCl_2_), bound proteins were eluted with 2% SDS sample buffer (90°C, 10 min), resolved by SDS–PAGE (Bio-Rad) and transferred to PVDF membrane for western blot analysis using indicated antibodies (see Reporting summary) anti-B55 (CST, 2290 S, 1:1,000), anti-PP2Ac (Millipore, MABE1783, 1:1,000), anti-PP2Aa (Biolegend, 824901, 1:1000), anti-Strep (Genscript, A01732, 1:1000), goat anti-rat IgG (Thermo Scientific, SA5-10024, 1:3000) goat anti-rabbi IgG (Bio-Rad, 12005869, 1:3,000) and goat anti-mouse IgG (Bio-Rad, 12004158, 1:3,000). Antibody fluorescence signals were captured using a ChemiDoc MP Imaging System (Image Lab Touch Software 2.4; Bio-Rad) and band intensities were quantified using ImageJ 1.53t. Uncropped blots are provided in Source data folder.

### EGFP_Nanobody protein expression, purification, and immobilization onto agarose beads

For expression, pOPIN_EGFP_Nanobody plasmid DNA (gift from M. Bollen, KU Leuven) was transformed into *E. coli* BL21 (DE3) cells (Agilent). Freshly transformed cells were grown at 37°C in LB broth containing carbenicillin antibiotics (50 µg/ml) until they reached an optical density (OD_600_) of ∼0.8. Protein expression was induced by addition of 0.5 mM β-D-thiogalactopyranoside (IPTG) to the culture medium, and cultures were allowed to grow overnight (18-20 hours, 250 rpm shaking) at 18°C. Cells were harvested by centrifugation (8000 *x*g, 15 min, 4°C) and stored at - 80°C until purification. Cell pellets expressing EGFP_Nanobody were resuspended in ice-cold lysis buffer (50 mM Tris pH 8.0, 500 mM NaCl, 5 mM Imidazole, 0.1% Triton X-100, EDTA-free protease inhibitor tablet [ThermoFisher]), lysed by high-pressure cell homogenization (Avestin Emulsiflex C3). Cell debris was pelleted by centrifugation (42,000 x*g*, 45 min, 4°C), and the supernatant was filtered with 0.22 µm syringe filters. The proteins were loaded onto a HisTrap HP column (Cytiva) pre-equilibrated with Buffer A (50 mM Tris pH 8.0, 500 mM NaCl, 5 mM imidazole) and eluted using a linear gradient (0-60% B) with buffer B (50 mM Tris pH 8.0, 500 mM NaCl and 500 mM imidazole). Fractions containing the protein were pooled, concentrated, and further purified at room temperature using size exclusion chromatography (SEC; Superdex 75 26/60 [Cytiva]) in PBS pH 7.5 buffer. Purified and concentrated EGFP_Nanobody protein was immobilized onto agarose beads (20 mg protein per column) using AminoLink Plus Immobilization Kit (ThermoFisher), following manufacturer’s instructions in PBS pH 7.5 coupling buffer.

### B55_LL_ purification

Expi293F cell pellets expressing EGFP_B55_LL_ were resuspended in ice-cold lysis buffer (20 mM Tris pH 8.0, 500 mM NaCl, 0.5 mM TCEP, 0.1% Triton X-100, EDTA-free protease inhibitor tablet [ThermoFisher]), lysed by high-pressure cell homogenization (Avestin Emulsiflex C3). Cell debris was pelleted by centrifugation (42,000 x*g*, 45 min, 4°C), and the supernatant was filtered with 0.22 µm syringe filters (Millipore). Lysates were mixed with GFP-nanobody-coupled agarose beads (see preparation in methods), pre-equilibrated with Wash Buffer 1 (20 mM Tris pH 8.0, 500 mM NaCl and 0.5 mM TCEP) and slowly rocked at 4°C for 2 hours. After two hours, lysate/beads mixture loaded onto gravity columns, the flow through (FT1) was collected and the column was washed 3 times with 25 mL of Wash Buffer (wash 1-3). The GFP-B55 resin was resuspended in Wash Buffer 2 (20 mM Tris pH 8.0, 250 mM NaCl and 0.5 mM TCEP) and TEV was added for on-column cleavage rocking overnight at 4°C. The flow through was again collected (FT2) and the resin was washed with 20 mL of Wash Buffer 2 (wash 4) and 2x 20 mL with the Wash Buffer 1 (wash 5 and 6). The flow through 2 (FT2) and washes 4-6 were collected, diluted to ∼100 mM salt concentration (with 0 mM NaCl Wash buffer), and loaded onto QTrap HP column (Cytiva) for further purification. The proteins were eluted with a 100 mM – 1 M salt gradient (Buffer A: 20 mM Tris pH 8.0, 100 mM NaCl, 0.5 mM TCEP; Buffer B: 20 mM Tris pH 8.0, 1 M NaCl, 0.5 mM TCEP). B55_LL_ was concentrated and further purified using SEC (Superdex 200 26/60 [Cytiva]) in either p107 or EYA3 NMR buffer or assay buffer (20 mM Tris pH 8.0, 150 mM NaCl, 0.5 mM TCEP).

### PP2A:B55 complex purification

*Expi293F* cell pellets expressing StrepII_PP2Ac and EGFP_B55 constructs were resuspended in ice-cold lysis buffer (20 mM Tris pH 8.0, 500 mM NaCl, 0.5 mM TCEP, 1 mM MnCl_2_, 0.1% Triton X-100, EDTA-free protease inhibitor tablet [ThermoFisher]), lysed by high-pressure cell homogenization (Avestin Emulsiflex C3). Purified PP2Aa was added to the cell lysate. Cell debris was pelleted by centrifugation (42,000 x*g*, 45 min, 4°C), and the supernatant was filtered with 0.22 µm syringe filters (Millipore). Lysates were loaded onto a GFP-Nanobody-coupled agarose bead (see preparation in methods) column, pre-equilibrated with Wash Buffer 1 (20 mM Tris pH 8.0, 500 mM NaCl, 1 mM MnCl_2_ and 0.5 mM TCEP) and slowly rocked at 4°C for 2 hours. After two hours, the flow through (FT1) was collected and the column was washed 3 times with 25 mL of Wash Buffer (wash 1-3). The GFP-B55 resin was resuspended in 20 mM Tris pH 8.0, 250 mM NaCl, 1 mM MnCl_2_ and 0.5 mM TCEP, and TEV was added for on-column cleavage rocking overnight at 4°C. The flow through was again collected (FT2) and the resin was washed with 20 mL of Wash Buffer 2 (20 mM Tris pH 8.0, 250 mM NaCl, 1 mM MnCl_2_ and 0.5 mM TCEP) (wash 4) and 2x 20 mL with the Wash Buffer 1 (wash 5 and 6). The flow through 2 (FT2) and washes 4-6 were collected, diluted to ∼100 mM salt concentration (with 0 mM NaCl Wash buffer), and loaded onto QTrap HP column (Cytiva) for further purification. The proteins were eluted with a 100 mM – 1 M salt gradient (Buffer A: 20 mM Tris pH 8.0, 100 mM NaCl, 1 mM MnCl_2_ and 0.5 mM TCEP; Buffer B: 20 mM Tris pH 8.0, 1 M NaCl, 1 mM MnCl_2_ and 0.5 mM TCEP). PP2A:B55 complex and B55 fractions were pooled, concentrated and further purified using SEC (Superdex 200 26/60 [Cytiva]) in either p107 or Eya3 NMR buffer or assay buffer (20 mM Tris pH 8.0, 150 mM NaCl, 1 mM MnCl_2_ and 0.5 mM TCEP).

### Cryo-EM sample preparation

The PP2A:B55-p107_612-687_ complex was prepared by purifying PP2A:B55 and incubating it with a 1.5 molar ratio of PP2A:B55-to-p107_612-687_ at a total concentration of 2.4 mg/ml. The PP2A:B55-Eya3_62-108EE_ complex was prepared by purifying PP2A:B55 and incubating it with a 1.56 molar ratio of PP2A:B55-to-Eya3_62-108EE_ at a total concentration of 2.4 mg/ml (**Extended Data Fig. 3**). Immediately prior to blotting and vitrification (Thermo Fisher Vitrobot MK IV, 18°C, 100% relative humidity), CHAPSO (3-([3-Cholamidopropyl]dimethylammonio)-2-hydroxy-1-propanesulfonate) was added to a final concentration of 0.125% (w/v) for PP2A:B55-p107_612-687_ and 0.1% (w/v) for PP2A:B55-Eya3_62-108EE_. 3.5 μL of the sample was applied to a freshly glow discharged UltAuFoil 1.2/1.3 300 mesh grid (SPT Labtech), blotted for 5 s and plunged into liquid ethane.

### Cryo-EM data acquisition and processing. PP2A:B55-p107_612-687_

Imaging was performed at the Pacific Northwest Cryo-EM Center (PNCC) using a Thermo Fisher Titan Krios G3i (fringe-free) operating at an accelerating voltage of 300 keV equipped with a Gatan BioQuantum energy filter and K3 camera operated in Super-res Counting mode. Acquisition and imaging parameters are given in **Table 1**. All data processing steps were performed using CryoSPARC4.4.1^45^ and are summarized in **Extended Data Fig. 4**. Super resolution movies were imported to CryoSPARC4.4.1 and motion corrected using patch motion correction. Initial contrast transfer function (CTF) parameters were then calculated using the patch CTF estimation. Micrographs were filtered to remove outliers in motion correction and/or CTF estimation results and screened manually to remove micrographs with significant non-vitreous ice contamination. 2D class averages of particles picked by blob picker from 4687 micrographs were used as templates for the template picker. Picks were subjected to 2D and 3D classification and a subset used to train a Topaz^46^ model, which was then used to pick 2.8 million particles. Multiple rounds of 2D and 3D classifications in CryoSPARC were used to remove junk particles from the initial picks, resulting in selected particles from classes showing clear secondary structure and representing the full complex. Resolution in both datasets was then further improved by cycles of CTF parameter refinement and particle polishing. A final round of 3D classification was used to identify particles with well-resolved density for p107. The final map (non-uniform refinement, CryoSPARC) was refined from 195,950 particles to a resolution of 2.58 Å. *PP2A:B55-EYA3_62-108EE_*: Imaging was performed at the NIH National Center for CryoEM Access and Training (NCCAT) using a Thermo Fisher Titan Krios G4 (fringe-free) operating at an accelerating voltage of 300 keV equipped with a Falcon 4i direct detector. Acquisition and imaging parameters are given in **Table 1**. All data processing steps were performed using CryoSPARC4.4.1 and are summarized in **Extended Data Fig. 5**. EER movies were imported to CryoSPARC4.4.1 and motion corrected using patch motion correction. Initial contrast transfer function (CTF) parameters were then calculated using the patch CTF estimation. Micrographs were filtered to remove outliers in motion correction and/or CTF estimation results and screened manually to remove micrographs with significant non-vitreous ice contamination. 2D class averages of particles picked by blob picker from 4084 micrographs were used as templates for the template picker. Picks were subjected to 2D and 3D classification and a subset used to train a Topaz model, which was then used to pick 874,057 particles. Multiple rounds of 2D and 3D classifications in CryoSPARC were used to remove junk particles from the initial picks, resulting in selected particles from classes showing clear secondary structure and representing the full complex. Resolution in both datasets was then further improved by cycles of CTF parameter refinement and particle polishing. A final round of 3D classification was used to identify particles with well-resolved density for Eya3. The final map (non-uniform refinement, CryoSPARC) was refined from 107,919 particles to a resolution of 2.7 Å. All global map resolutions reported in this work were calculated by the gold-standard half-maps FSC=0.143 metric as reported in CryoSPARC. The structures were visualized using PyMOL (PyMOL Molecular Graphics System, Schroedinger, LLC) or UCSF ChimeraX (v1.2.5)^47^.

### Cryo-EM model building

All models were built and refined by iterating between manual rebuilding and refinement in Coot^48^ and automated global real-space refinement in Phenix^49^ using the unsharpened maps. For both complexes, PP2Aa, B55 and PP2Ac from the PP2A:B55-FAM122A complex (PDBID 8SO0) were docked into the maps using ChimeraX. Density corresponding to p107 was clearly helical while density corresponding to Eya3 was extended, with the strongest density at the bottom of the B55 platform showing a clear π-stacking interaction with H179_B55_. p107 and Eya3 residues identified to interact with B55 by NMR were readily modeled into the density. Model geometry and map-model validation metrics are given in **Table 1**.

### DiFMUP fluorescence intensity assay for PP2A:B55 IC_50_ measurements

DiFMUP based IC_50_ assays were conducted in 384 well plates (Corning, cat# 4411). For p107_612-687_ and Eya3_62-108EE_ IC_50_ assays, PP2A:B55 holoenzyme in Enzyme buffer (30 mM HEPES pH 7.0, 150 mM NaCl, 1 mM MnCl_2_, 1 mM DTT, 0.01% triton X-100, 0.1 mg/ml BSA) was pre-incubated with various concentrations of p107_612-687_ and Eya3_62-108EE_ variants for 30 min at room temperature. The reaction was started by adding DiFMUP (final concentration 50 μM) into the PP2A:B55-p107_612-687_ and PP2A:B55-Eya3_62-108EE_ enzymatic reaction (final concentration of PP2A:B55 holoenzyme at 1 nM) and then incubated at 30°C for 30 min. End-point reads (ex=360 nm, em=450 nm) were taken on a CLARIOstarPlus (BMG LABTECH) plate reader (using reader control software version 5.7 R2) after the reaction was stopped by the addition of 300 mM potassium phosphate (pH 10). The experiments were independently technically repeated ≥ three times (each reaction was made in n = 3 to 6) and the averaged IC_50_ and standard deviation (SD) values were reported. The data was evaluated using GraphPad Prism 10.2.2.

### NMR data collection

All NMR data were collected on either a Bruker Avance Neo 600 MHz or 800 MHz NMR spectrometer equipped with TCI HCN z-gradient cryoprobe at 283 K. (^15^N,^13^C)-labeled Eya3 (170 µM) was prepared in Eya3 NMR buffer with 5-10% (v/v) D_2_O added immediately prior to data acquisition. The sequence-specific backbone assignment was determined by recording a suite of heteronuclear NMR spectra: 2D [^1^H,^15^N] HSQC, 3D HNCA, 3D HN(CO)CA, 3D HNCACB, 3D CBCA(CO)NH, 3D HNCO, and 3D HN(CA)CO. Spectra were processed in Topspin (Bruker Topspin 4.1.3) and referenced to internal DSS.

### Sequence-specific backbone assignment, chemical shift index and chemical shift perturbation

Peak picking and sequence-specific backbone assignment were performed using the program CARA 1.9.1 (http://www.cara.nmr.ch). Chemical shift index (CSI) calculations of p107 and Eya3 were performed using both Cα and Cβ chemical shifts for each assigned amino acid, omitting glycine, against the RefDB database^50^. Secondary structure propensity (SSP) scores were calculated using a weighted average of seven residues to minimize contributions from chemical shifts of residues that are poor measures of secondary structure. Chemical shift differences (Δδ) were calculated using the following equation:

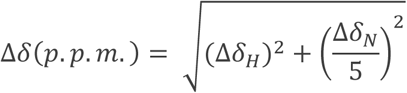

### NMR interaction studies of p107/EYA3 with PP2A:B55 and B55_LL_

All NMR interaction data of p107/Eya3 with either PP2A:B55 or B55_LL_ was recorded using either a Bruker Neo 600 MHz or 800 MHz NMR spectrometer equipped with a HCN TCI active z-gradient cryoprobe at 283 K. All NMR measurements of p107 or Eya3 were recorded using ^15^N-labeled protein in NMR buffer and 90% H_2_O/10% D_2_O. For each interaction, an excess of unlabeled B55_LL_ or PP2A:B55 complex (min 25% surplus ratio) was added to ^15^N-labeled p107/Eya3 and incubated on ice for 10 minutes before the 2D [^1^H,^15^N] HSQC spectrum was collected. p107/Eya3 concentrations ranged from 2.5-6 μM. NMR data were processed using nmrPipe^51^ and the intensity data were analyzed in Poky.^52^ The p107 intensity data was normalized to its C-terminal residue peak and the difference between the free 2D [^1^H,^15^N] HSQC spectrum of p107 was compared to its respective peak, if present, on the 2D [^1^H,^15^N] HSQC spectrum of p107 in complex with B55_LL_ or PP2A:B55. The Eya3 intensity data was normalized to the average of the last two c-terminal residue (EE) peaks and the difference between the free 2D [^1^H,^15^N] HSQC spectrum of Eya3 was compared to its respective peak, if present, on the 2D [^1^H,^15^N] HSQC spectrum of Eya3 in complex with B55_LL_ or PP2A:B55. Any overlapping peaks were omitted for this analysis. CCPNMR was used for spectra overlays and figures^53^.

### NMR dephosphorylation

All dephosphorylation data was recorded using a Bruker Neo 600 MHz NMR spectrometer equipped with a HCN TCI active z-gradient cryoprobe at 283 K. All measurements were recorded using ^15^N-labeled p107 in p107 NMR buffer and 90% H_2_O/10% D_2_O. A reference 2D [^1^H,^15^N] HSQC was recorded. Unlabeled active PP2A:B55 was added to ^15^N-labeled p107 in 1:500 ratio. A 2D [^1^H,^15^N] HSQC spectra was measured every 28 minutes to monitor dephosphorylation. NMR data were processed using nmrPipe^51^ and the disappearing intensities of the phosphorylated peaks, and the reappearing intensity of the unphosphorylated peaks were analyzed with Poky^52^.

## Data Availability

The NMR data generated in this study have been deposited in the BioMagResBank database under accession code BMRB 52492 (https://bmrb.io/data_library/summary/?bmrbId=52492) (Eya3). The atomic coordinates and structure factors for the PP2A:B55-p107 complex have been deposited in the PDB database under accession code 9C6B (10.2210/pdb9C6B/pdb) and EMDB code EMD-45243. The atomic coordinates and structure factors for PP2A:B55-Eya3 complex have been deposited in the PDB database under accession code 9C7T (10.2210/pdb9C7T/pdb) and EMDB code EMD-45292. All IC_50_ and pull-down data generated in this study are provided in the Supplementary Information and/or Source Data file, which is available at Figshare (doi.org/10.6084/m9.figshare.26278654.v1).

## Acknowledgements

We thank Dr. Steven Blacklow for sharing the PDB file of the PP2A:B55-IRE5 structure in advance of publication. This work was supported by grant 1R01GM144483 from the National Institute of General Medicine and 1R01NS124666 from the National Institute of Neurological Disorders and Stroke to WP and grant 1R01GM144379 from the National Institute of General Medicine to RP. Cryo-EM grid screening was performed on a Tundra 100 keV cryo-EM instrument supported by NIH S10OD032156. Portions of this research were (1) supported by NIH grant U24GM129547 and performed at the PNCC at OHSU and accessed through EMSL (grid.436923.9), a DOE Office of Science User Facility sponsored by the Office of Biological and Environmental Research and (2) performed at the National Center for CryoEM Access and Training (NCCAT) and the Simons Electron Microscopy Center located at the New York Structural Biology Center, supported by the NIH Common Fund Transformative High Resolution Cryo-Electron Microscopy program (U24 GM129539 and R24GM154192) and by grants from the Simons Foundation (SF349247) and NY State Assembly.

## Author Contributions

RP, WP, SP and RJG developed the concept. SK and RJG expressed and purified all proteins. RJG and WP performed and analyzed NMR experiments. SK and RP determined the cryo-EM structures. SP, RJG performed pulldown and IC_50_ work. RP, WP, SP and RJG wrote the manuscript.

## Competing Interests Statement

The authors declare no competing interests. The funders had no role in study design, data collection and analysis, decision to publish, or preparation of the manuscript.

## Notes

### Competing Interest Statement

The authors have declared no competing interest.

## REFERENCES

1. Brautigan, D. L. & Shenolikar, S. Protein Serine/Threonine Phosphatases: Keys to Unlocking Regulators and Substrates. Annu. Rev. Biochem. 87, 921–964 (2018).

2. Johnson, J. L. et al. An atlas of substrate specificities for the human serine/threonine kinome. Nature 613, 759–766 (2023).

3. Nguyen, H. & Kettenbach, A. N. Substrate and phosphorylation site selection by phosphoprotein phosphatases. Trends Biochem Sci 48, 713–725 (2023).

4. Kokot, T. & Köhn, M. Emerging insights into serine/threonine-specific phosphoprotein phosphatase function and selectivity. J Cell Sci 135, jcs259618 (2022).

5. Hendus-Altenburger, R. et al. Molecular basis for the binding and selective dephosphorylation of Na+/H+ exchanger 1 by calcineurin. Nat Commun 10, 3489 (2019).

6. Grigoriu, S. et al. The molecular mechanism of substrate engagement and immunosuppressant inhibition of calcineurin. PLoS Biol 11, e1001492 (2013).

7. Wang, X., Bajaj, R., Bollen, M., Peti, W. & Page, R. Expanding the PP2A Interactome by Defining a B56-Specific SLiM. Structure 24, 2174–2181 (2016).

8. Ragusa, M. J. et al. Spinophilin directs protein phosphatase 1 specificity by blocking substrate binding sites. Nat Struct Mol Biol 17, 459–464 (2010).

9. Ueki, Y. et al. A Consensus Binding Motif for the PP4 Protein Phosphatase. Mol Cell 76, 953–964.e6 (2019).

10. Hertz, E. P. T. et al. A Conserved Motif Provides Binding Specificity to the PP2A-B56 Phosphatase. Mol. Cell 63, 686–695 (2016).

11. Peti, W., Nairn, A. C. & Page, R. Structural basis for protein phosphatase 1 regulation and specificity. FEBS J 280, 596–611 (2013).

12. O’Connell, N. et al. The molecular basis for substrate specificity of the nuclear NIPP1:PP1 holoenzyme. Structure 20, 1746–1756 (2012).

13. Choy, M. S. et al. Understanding the antagonism of retinoblastoma protein dephosphorylation by PNUTS provides insights into the PP1 regulatory code. Proc Natl Acad Sci U S A 111, 4097–4102 (2014).

14. Fedoryshchak, R. O. et al. Molecular basis for substrate specificity of the Phactr1/PP1 phosphatase holoenzyme. Elife 9, e61509 (2020).

15. Hurley, T. D. et al. Structural basis for regulation of protein phosphatase 1 by inhibitor-2. J. Biol. Chem. 282, 28874–28883 (2007).

16. Aramburu, J. et al. Selective inhibition of NFAT activation by a peptide spanning the calcineurin targeting site of NFAT. Mol Cell 1, 627–637 (1998).

17. Hubbard, M. J. & Cohen, P. On target with a new mechanism for the regulation of protein phosphorylation. Trends Biochem Sci 18, 172–177 (1993).

18. Kruse, T. et al. The Ebola Virus Nucleoprotein Recruits the Host PP2A-B56 Phosphatase to Activate Transcriptional Support Activity of VP30. Mol. Cell 69, 136–145.e6 (2018).

19. Ammosova, T. et al. Nuclear targeting of protein phosphatase-1 by HIV-1 Tat protein. J Biol Chem 280, 36364–36371 (2005).

20. Shi, Y. Serine/threonine phosphatases: mechanism through structure. Cell 139, 468–484 (2009).

21. Yoo, S. J.-S., Boylan, J. M., Brautigan, D. L. & Gruppuso, P. A. Subunit composition and developmental regulation of hepatic protein phosphatase 2A (PP2A). Arch Biochem Biophys 461, 186–193 (2007).

22. Tung, H. Y., Resink, T. J., Hemmings, B. A., Shenolikar, S. & Cohen, P. The catalytic subunits of protein phosphatase-1 and protein phosphatase 2A are distinct gene products. Eur J Biochem 138, 635–641 (1984).

23. Wang, X. et al. A dynamic charge-charge interaction modulates PP2A:B56 substrate recruitment. Elife 9, (2020).

24. Tung, H. Y., Alemany, S. & Cohen, P. The protein phosphatases involved in cellular regulation. 2. Purification, subunit structure and properties of protein phosphatases-2A0, 2A1, and 2A2 from rabbit skeletal muscle. Eur J Biochem 148, 253–263 (1985).

25. Xu, Y., Chen, Y., Zhang, P., Jeffrey, P. D. & Shi, Y. Structure of a protein phosphatase 2A holoenzyme: insights into B55-mediated Tau dephosphorylation. Mol. Cell 31, 873–885 (2008).

26. Padi, S. K. R. et al. Cryo-EM structures of PP2A:B55-FAM122A and PP2A:B55-ARPP19. Nature 625, 195–203 (2024).

27. Cao, R. et al. Molecular Mechanism of PP2A/B55α Inhibition by IER5. bioRxiv 2023.08.29.555174 (2023) doi:10.1101/2023.08.29.555174.

28. Zhu, L. et al. Inhibition of cell proliferation by p107, a relative of the retinoblastoma protein. Genes Dev 7, 1111–1125 (1993).

29. Ginsberg, D. et al. E2F-4, a new member of the E2F transcription factor family, interacts with p107. Genes Dev 8, 2665–2679 (1994).

30. Fowle, H. et al. PP2A/B55α substrate recruitment as defined by the retinoblastoma-related protein p107. Elife 10, e63181 (2021).

31. Jayadeva, G. et al. B55alpha PP2A holoenzymes modulate the phosphorylation status of the retinoblastoma-related protein p107 and its activation. J Biol Chem 285, 29863–29873 (2010).

32. Jemc, J. & Rebay, I. The eyes absent family of phosphotyrosine phosphatases: properties and roles in developmental regulation of transcription. Annu Rev Biochem 76, 513–538 (2007).

33. Bui, Q. T., Zimmerman, J. E., Liu, H. & Bonini, N. M. Molecular analysis of Drosophila eyes absent mutants reveals features of the conserved Eya domain. Genetics 155, 709–720 (2000).

34. Jemc, J. & Rebay, I. Identification of transcriptional targets of the dual-function transcription factor/phosphatase eyes absent. Dev Biol 310, 416–429 (2007).

35. Alderman, C. et al. Biochemical characterization of the Eya and PP2A-B55α interaction. J Biol Chem 107408 (2024) doi:10.1016/j.jbc.2024.107408.

36. Sano, T. & Nagata, S. Characterization of the threonine-phosphatase of mouse eyes absent 3. FEBS Lett 585, 2714–2719 (2011).

37. Zhang, L. et al. Eya3 partners with PP2A to induce c-Myc stabilization and tumor progression. Nat Commun 9, 1047 (2018).

38. Bertran, M. T. et al. ASPP proteins discriminate between PP1 catalytic subunits through their SH3 domain and the PP1 C-tail. Nat Commun 10, 771 (2019).

39. Hornbeck, P. V., Chabra, I., Kornhauser, J. M., Skrzypek, E. & Zhang, B. PhosphoSite: A bioinformatics resource dedicated to physiological protein phosphorylation. Proteomics 4, 1551–1561 (2004).

40. Lowe, E. D. et al. Specificity determinants of recruitment peptides bound to phospho-CDK2/cyclin A. Biochemistry 41, 15625–15634 (2002).

41. Dyson, H. J. & Wright, P. E. NMR illuminates intrinsic disorder. Curr Opin Struct Biol 70, 44– 52 (2021).

42. Fuxreiter, M. Context-dependent, fuzzy protein interactions: Towards sequence-based insights. Curr Opin Struct Biol 87, 102834 (2024).

43. Fuxreiter, M. Electrostatics tunes protein interactions to context. Proc Natl Acad Sci U S A 119, e2209201119 (2022).

44. Fuxreiter, M. Fuzziness: linking regulation to protein dynamics. Mol Biosyst 8, 168–177 (2012).

45. McSweeney, D. M., McSweeney, S. M. & Liu, Q. A self-supervised workflow for particle picking in cryo-EM. IUCrJ 7, 719–727 (2020).

46. Bepler, T. et al. Positive-unlabeled convolutional neural networks for particle picking in cryo-electron micrographs. Nat Methods 16, 1153–1160 (2019).

47. Pettersen, E. F. et al. UCSF ChimeraX: Structure visualization for researchers, educators, and developers. Protein Sci 30, 70–82 (2021).

48. Emsley, P. & Cowtan, K. Coot: model-building tools for molecular graphics. Acta Crystallogr. D Biol. Crystallogr. 60, 2126–2132 (2004).

49. Adams, P. D. et al. PHENIX: a comprehensive Python-based system for macromolecular structure solution. Acta Crystallogr. D Biol. Crystallogr. 66, 213–221 (2010).

50. Zhang, H., Neal, S. & Wishart, D. S. RefDB: a database of uniformly referenced protein chemical shifts. J Biomol NMR 25, 173–195 (2003).

51. Delaglio, F. et al. NMRPipe: A multidimensional spectral processing system based on UNIX pipes. J Biomol NMR 6, 277–293 (1995).

52. Lee, W., Rahimi, M., Lee, Y. & Chiu, A. POKY: a software suite for multidimensional NMR and 3D structure calculation of biomolecules. Bioinformatics 37, 3041–3042 (2021).

53. Skinner, S. P. et al. CcpNmr AnalysisAssign: a flexible platform for integrated NMR analysis. J Biomol NMR 66, 111–124 (2016).

54. Potter, S. C. et al. HMMER web server: 2018 update. Nucleic Acids Res 46, W200–W204 (2018).

