## Supplemental Data for "Cryo-EM structures of PP2A:B55-Eya3 and PP2A:B55-p107 define PP2A:B55 substrate recruitment"

**Table of Contents.**

**Extended Data Table 1.** Inhibition ($IC_{50}$) of PP2A:B55 by p107 and Eya3

**Extended Data Figure 1.** p107 binds PP2A:B55

**Extended Data Figure 2.** Characterization of p107 and Eya3 in the free state

**Extended Data Figure 3.** PP2A:B55-p107 and PP2A:B55-Eya3 grid preparation

**Extended Data Figure 4.** Cryo-EM image processing workflow and maps for PP2A:B55-p107

**Extended Data Figure 5.** Cryo-EM image processing workflow and maps for PP2A:B55-Eya3

**Extended Data Figure 6.** PP2A:B55 preferentially dephosphorylates p107 CDK2/Cyclin A2 phosphosite *p*S640, full spectra

**Extended Data Figure 7.** p107, *p*S615 and *p*S640 dephosphorylation by PP2A:B55

**Extended Data Table 1. Inhibition ($IC_{50}$) of PP2A:B55 by p107 and Eya3**

| Substrate/Regulator | $IC_{50}$ [nM] | n |
|---|---|---|
| p107 | 245 ± 27 | 3 |
| Eya3 | 1102 ± 84 | 3 |

All experiments were performed as experimental triplicates
mean ± s.d., n = number of independent measurements.

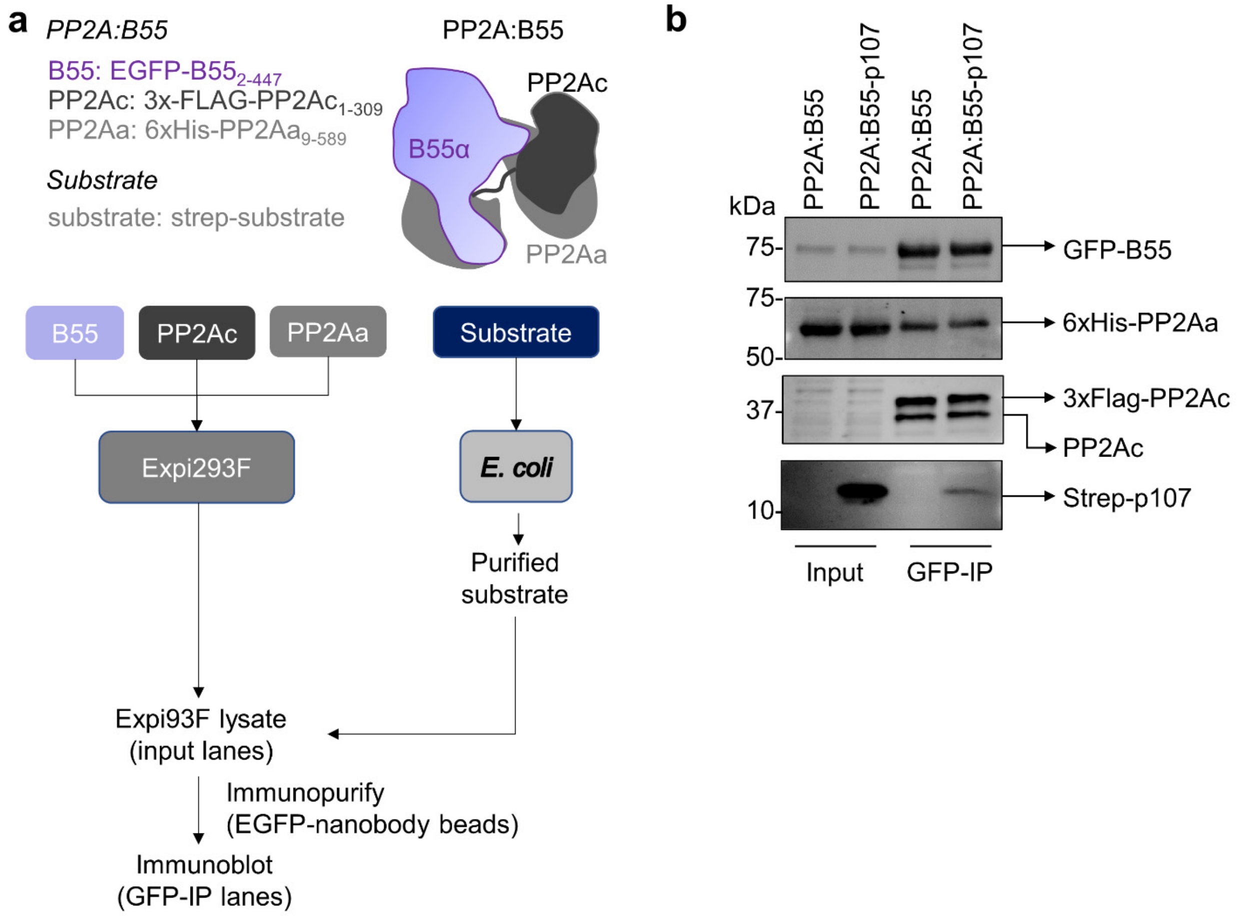


**Extended Data Figure 1. p107 binds PP2A:B55. a.** Pull-down assay schemes, with input and GFP-IP steps indicated. **b**. Expi293F cell lysates expressing GFP-B55, 3xFLAG-PP2Ac and 6xHis-PP2Aa were incubated with or without purified p107 (residues 612-687), PP2A:GFP-B55 complexes immunopurified using GFP-trap beads (EGFP nanobodies coupled to agarose resin) and isolated proteins detected by immunoblot (representative of 3 independent experiments).

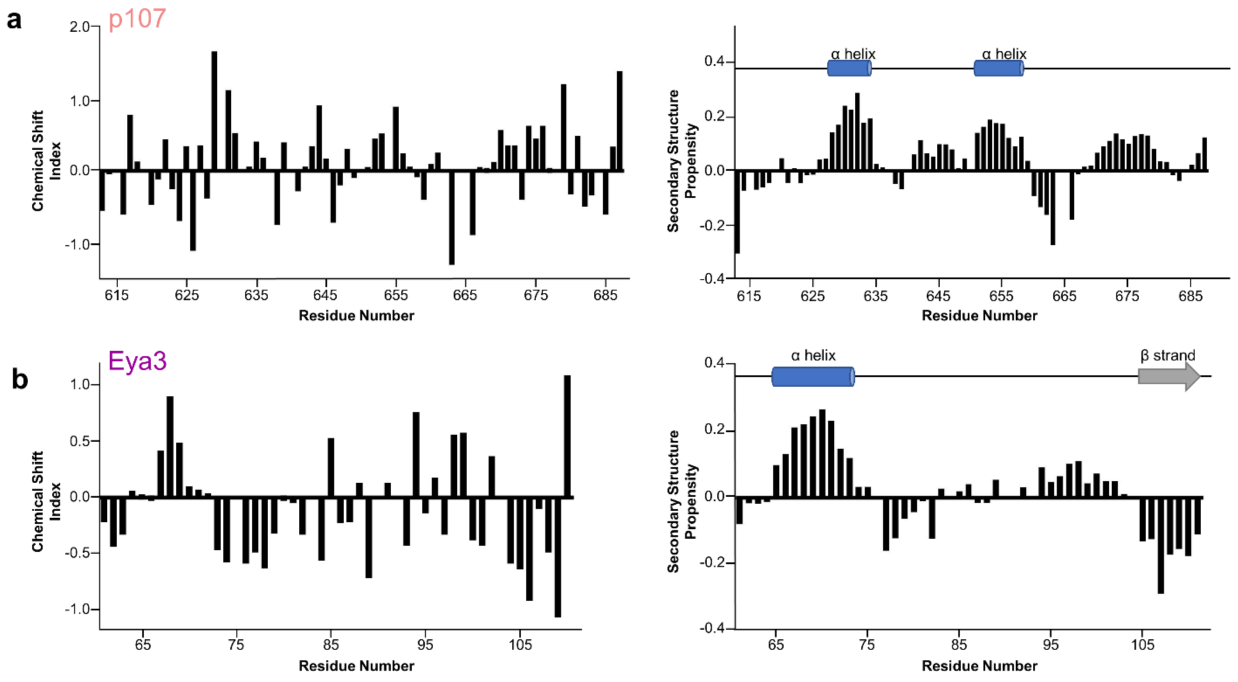


**Extended Data Figure 2. Characterization of p107 and Eya3 in the free state. a.** Chemical Shift Index (CSI) (left) and Secondary-structure propensity (SSP) (right) data for p107 plotted vs. residue numbers. (SSP > 0, α helix; SSP < 0, β strand). Cα and Cβ chemical shifts were used to create the CSI and SSP plots (RefDB database)[50]. Preferred secondary structure indicated above SSP data. **b.** CSI (left) and SSP (right) data for Eya3 plotted vs. residue numbers, same as in (a).

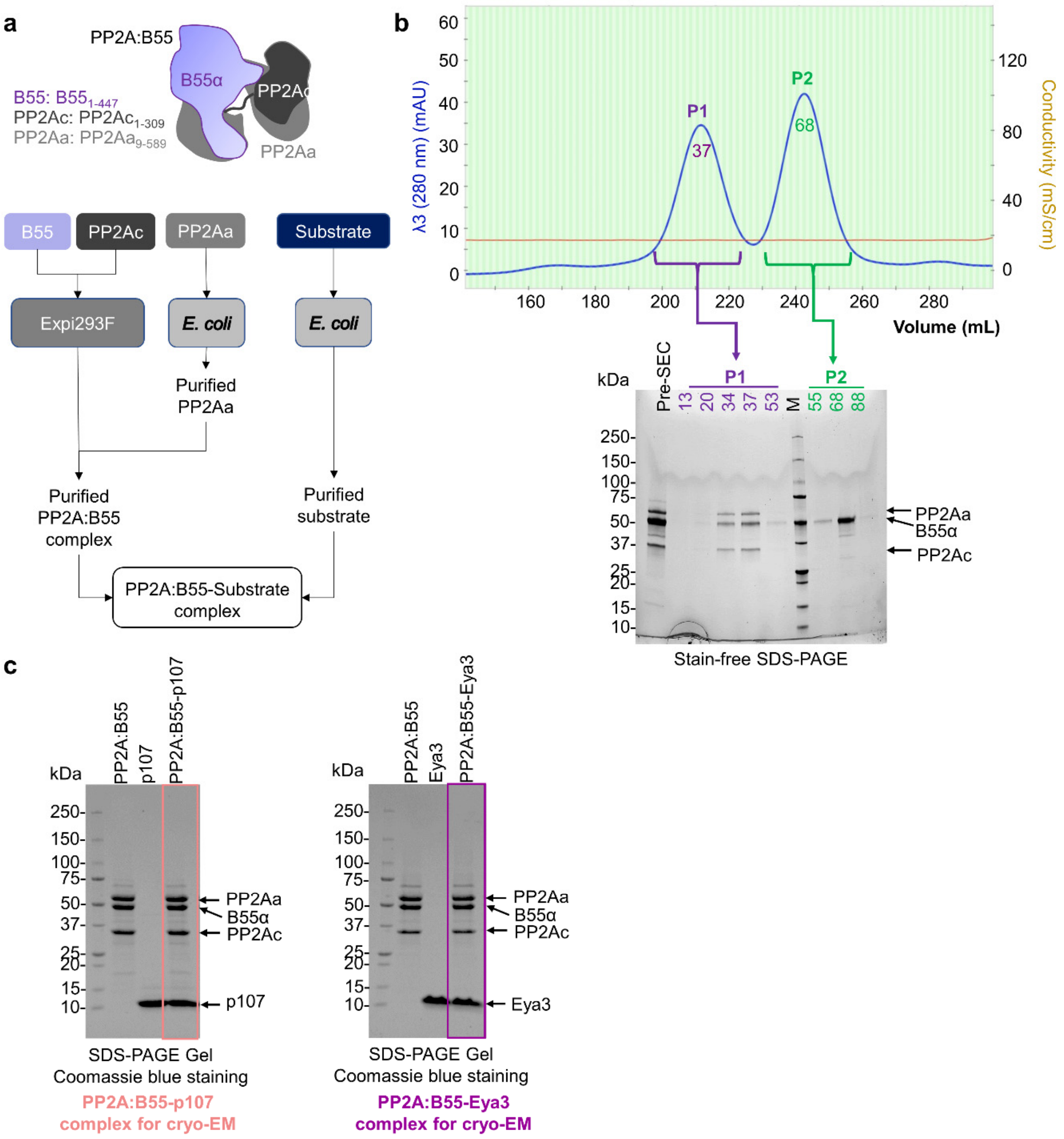


| Cryo-EM sample | Molar ratio | Protein Concentration | CHAPSO |
|---|---|---|---|
| PP2A:B55-p107 | 1:5 | 2.4 mg/mL | 0.125% |
| PP2a:B55-Eya3 | 1:5.6 | 2.4 mg/mL | 0.1% |

**Extended Data Figure 3. Purification of active, methylated PP2A:B55 from human cells. a.** Schematic of the production of PP2A:B55 and PP2A:B55-substrate complexes for structural and biophysical studies. **b.** Size exclusion chromatography of PP2A:B55, demonstrating the PP2A-B55 complex is stable and elutes as a single peak (Peak 1) with excess B55 eluting separately (peak 2), as shown in stain-free SDS-PAGE. **c.** SDS-PAGE of PP2A:B55-p107 and PP2A:B55-Eya3 samples (top), including protein and detergent concentrations (bottom), used for cryo-EM.

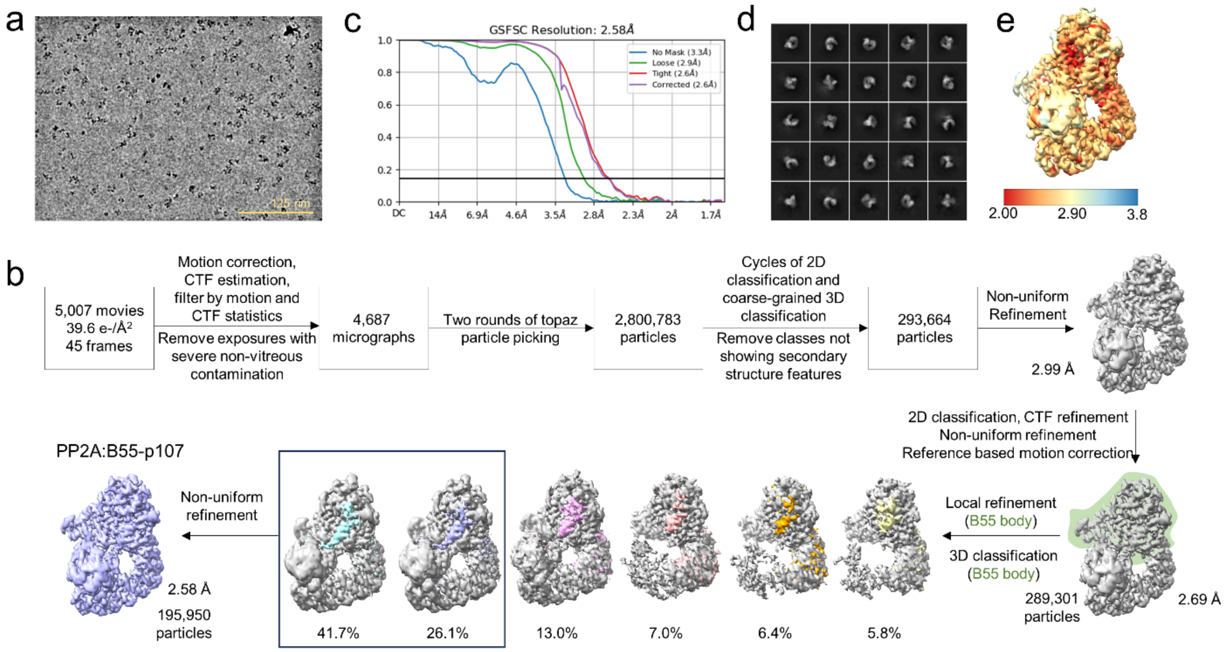


**Extended Data Figure 4. Cryo-EM image processing workflow and maps for PP2A:B55-p107. a.** Representative micrograph. **b**. Particle counts and reconstruction resolution are given at key junctions of the process (all resolutions reported are calculated by the gold standard half-maps FSC=0.143 criterion). **c**. The global resolution estimate from the masked Fourier Shell Correlation curve is 2.58 Å at FSC of 0.143. **d**. Reference-free 2D class averages generated from the 195,950 particles used in the final refinement. **e**. Cryo-EM map colored by local resolution.

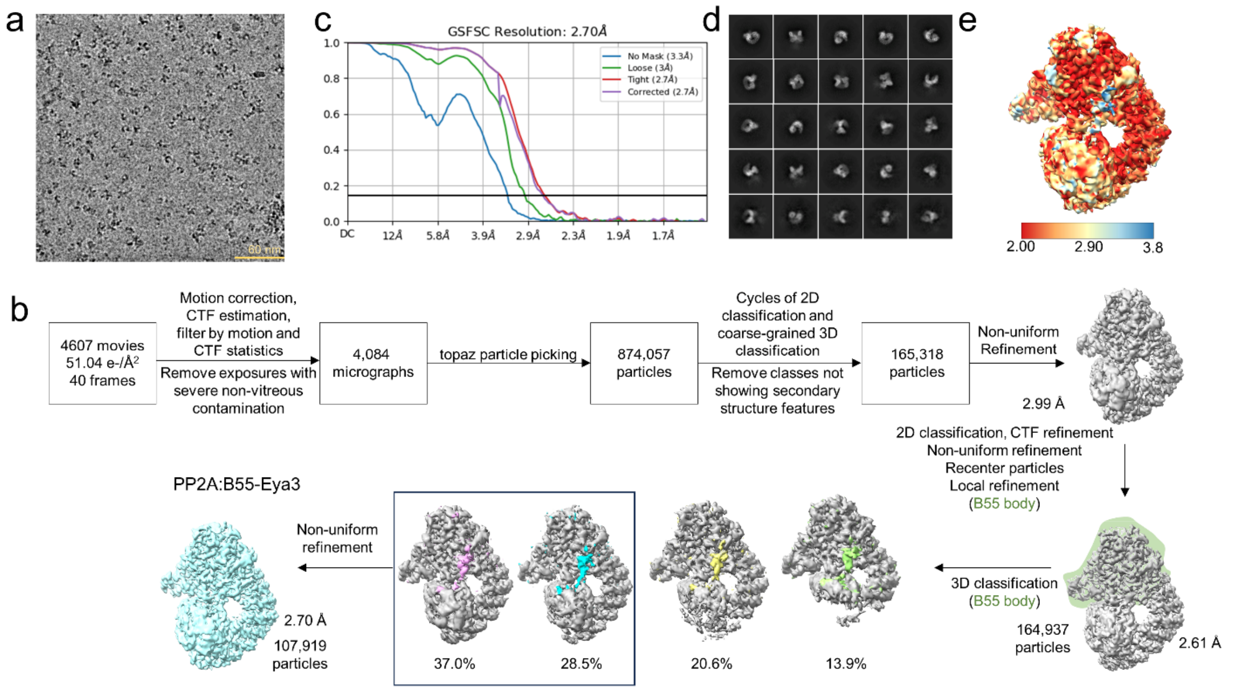


**Extended Data Figure 5. Cryo-EM image processing workflow and maps for PP2A:B55-Eya3. a.** Representative micrograph. **b**. Particle counts and reconstruction resolution are given at key junctions of the process (all resolutions reported are calculated by the gold standard half-maps FSC=0.143 criterion). **c**. The global resolution estimate from the masked Fourier Shell Correlation curve is 2.70 Å at FSC of 0.143. **d**. Reference-free 2D class averages generated from the 107,919 particles used in the final refinement. **e**. Cryo-EM map colored by local resolution.

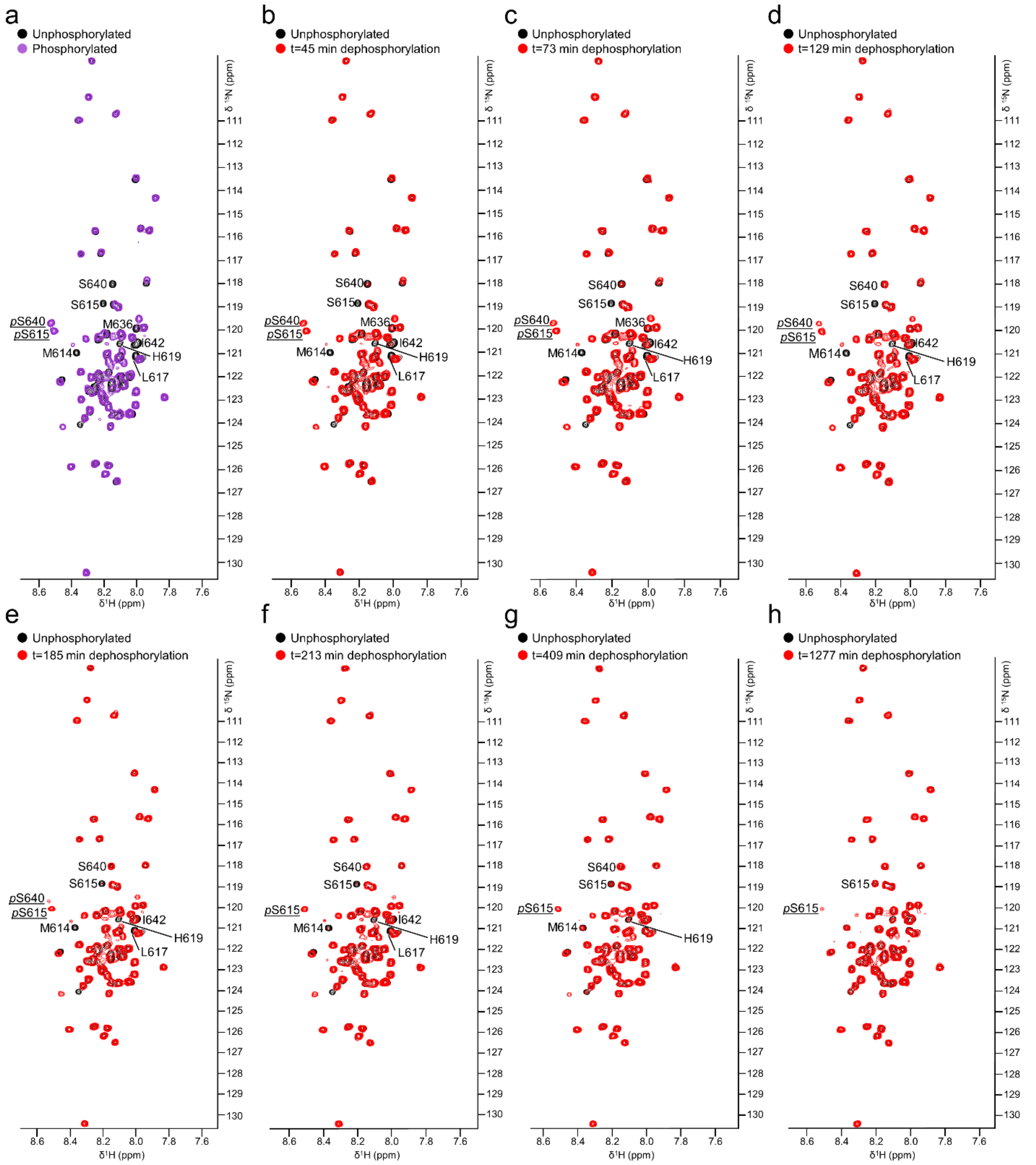


**Extended Data Figure 6. PP2A:B55 preferentially dephosphorylates p107 CDK2/Cyclin A2 phosphosite *p*S640, full spectra. a**. 2D [$^{1}$H,$^{15}$N] HSQC spectrum of $^{15}$N-labeled unphosphorylated p107 (black) overlaid with that of CDK2/Cyclin A2 phosphorylated $^{15}$N-labeled p107 (purple); *p*S615, *p*S640 and shifted peaks labeled. **b-h**. 2D [$^{1}$H,$^{15}$N] HSQC spectra of $^{15}$N-labeled unphosphorylated p107 (black) overlaid with that of CDK2/Cyclin A2 phosphorylated $^{15}$N-labeled p107 incubated with PP2A:B55 (red) for the following timepoints in minutes: 45 (b), 73 (c), 129 (d), 185 (e), 213 (f), 409 (g) and 1277 (h).

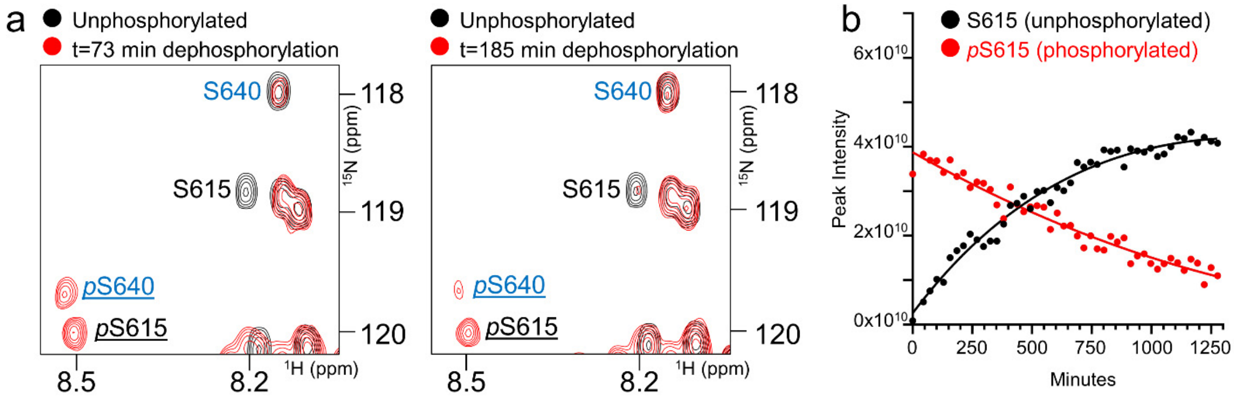


**Extended Data Figure 7. p107, *p*S615 and *p*S640 dephosphorylation by PP2A:B55. a.** 2D [$^1$H,$^{15}$N] HSQC spectrum of unphosphorylated $^{15}$N-labeled p107 (black) overlaid with that of CDK2/Cyclin A2 phosphorylated $^{15}$N-labeled p107 incubated with PP2A:B55 (red) for 73 minutes (left) and 185 minutes (right). **b.** Changes in peak intensities of $^{15}$N-labeled p107 residues *p*S615 (red) and S615 (black).
